# Mating-type locus rearrangement leads to shift from homothallism to heterothallism in *Citrus*-associated *Phyllosticta* species

**DOI:** 10.1101/2020.04.14.040725

**Authors:** Desirrê Alexia Lourenço Petters-Vandresen, Bruno Janoski Rossi, Johannes Z. Groenewald, Pedro W. Crous, Marcos Antonio Machado, Eva H. Stukenbrock, Chirlei Glienke

## Abstract

Currently, eight *Phyllosticta* species are known to be associated with *Citrus* hosts, incorporating endophytic and pathogenic lifestyles. As sexual reproduction is a key factor involved in host-interaction, it could be related to the differences in lifestyle. To evaluate this hypothesis, we characterized the mating-type loci of six *Citrus-*associated *Phyllosticta* species from whole genome assemblies. Mating-type genes are highly variable in their sequence content, but the genomic locations and organization of the mating-type loci are conserved. *Phyllosticta citriasiana, P. citribraziliensis* and *P. paracitricarpa* are heterothallic, and *P. citrichinaensis* was confirmed to be homothallic. In addition, the *P. citrichinaensis MAT1-2* idiomorph occurs in a separate location from the mating-type locus. Ancestral state reconstruction suggests that homothallism is the ancestral thallism state in *Phyllosticta*, with a shift to heterothallism in *Phyllosticta* species that are pathogenic to *Citrus*. Moreover, the homothallic strategies of *P. capitalensis* and *P. citrichinaensis* result from independent evolutionary events. As the pathogenic species *P. citriasiana, P. citricarpa* and *P. paracitricarpa* are heterothallic and incapable of selfing, disease management practices focused in preventing the occurrence of sexual reproduction could assist in the control of Citrus Black Spot and Citrus Tan Spot diseases. This study emphasizes the importance of studying *Citrus*-*Phyllosticta* interactions under evolutionary and genomic perspectives, as these approaches can provide valuable information about the association between *Phyllosticta* species and their hosts, and also serve as guidance for the improvement of disease management practices.

## 1 Introduction

Sexual reproduction plays a key role in the evolution of fungi and other eukaryotic organisms, by increasing genetic variation, purging deleterious mutations, and mediating the production of resistant sexual spores (Lee et al., 2010; Ni et al., 2011). The core features of sexual reproduction, such as ploidy changes, production of gametes via meiosis, cell-cell recognition and fusion, are conserved along all eukaryotic organisms (Heitman et al., 2013). Nevertheless, fungal species present diverse reproduction strategies to complete the sexual cycle (Ni et al., 2011). In some species, including *Coprinopsis cinerea* and *Schizophyllum commune*, sexual reproduction is regulated by multiple mating-types (Casselton and Kües, 2007); at the other extreme some species are self-fertile, such as *Aspergillus nidulans* (Kronstad, 2007).

Sexual reproduction in fungi can be classified in two main strategies, namely heterothallism and homothallism. In Pezizomycotina, the differences between heterothallic and homothallic strategies arise from a bipolar mating-system (Ni et al., 2011), in which a single locus (*MAT1*) regulates mating processes in strains of the same species. Instead of alleles, the *MAT1* locus has two highly divergent versions (*MAT1-1* and *MAT1-2* idiomorphs) comprised of non-orthologous genes, and both are required for the successful completion of sexual reproduction (Wilken et al., 2017). In heterothallic species, individuals harbour only one *MAT1* idiomorph in their genomes, never both (Wilken et al., 2017). Thus, two individuals of opposite and compatible mating-types are required for sexual reproduction. On the other hand, the homothallic species possess both *MAT1* idiomorphs in their genomes. Therefore, each individual can reproduce with any other individual from the same species, including itself, because individuals are self-fertile (Nagel et al., 2018).

The *MAT1-1* idiomorph is characterized by a gene (*MAT1-1-1*) encoding a protein with an alpha-box domain (CDD: cl07856, pfam04769), while the *MAT1-2* idiomorph comprises a gene (*MAT1-2-1*) encoding a protein with a high mobility group domain (HMG box, CDD: cd01389) (Bihon et al., 2014). These mating-type genes are key components in sexual mating, and are transcription factors that control the sexual development and the downstream-regulation of the expression of mating-type specific genes (Kim et al., 2015). In addition to *MAT1-1-1* and *MAT1-2-1*, additional genes may exist in each idiomorph, and their nomenclature follows the system as originally described (Turgeon and Yoder, 2000; Wilken et al., 2017).

Even though the *MAT1* idiomorphs are not orthologous and show a high degree of variation among species, alignments of the sequences flanking the mating-type locus exhibit a high level of similarity between related species (Nagel et al., 2018). Sequence comparisons among Ascomycete species revealed that *MAT1-1* and *MAT1-2* flanking regions mostly encode the genes *SLA2, APN2*/DNA lyase, as well as *APC5* and *COX13* (Wilken et al., 2017). These genes show a higher degree of sequence conservation than *MAT1* idiomorphs. The presence of these flanking genes along several ascomycete species suggests a high degree of positional conservation for the mating-type locus in the genomes (Wilken et al., 2017).

The identification and characterization of the *MAT1* locus and adjacent regions is the first step to study and investigate the reproductive strategies of Ascomycete species (Wilken et al., 2017). Traditionally this has been done using molecular approaches, such as cloning and hybridization with specific probes (Glass et al., 1988), PCR with degenerate primers (Kerényi et al., 2004) and gene-walking approaches (Groenewald et al., 2006). Recently, with the advance of whole genome sequencing, BLAST analyses combined with gene prediction tools have been widely employed to identify and characterize the *MAT1* locus in different fungal species (Amorim et al., 2017; Bihon et al., 2014; Guarnaccia et al., 2019; Nagel et al., 2018; Wang et al., 2016). For species in which the sexual structures have been described (Robinson and Natvig, 2019), characterization of the *MAT1* locus helps to determine if the species is homo- or heterothallic, information that is essential to design crossing experiments. Likewise, in species for which sexual structures are not described and for which experimental crossing is not possible, analyses of the *MAT* locus have been important understand the species’ life cycle and sexual reproduction strategies (Linde et al., 2003; Waalwijk et al., 2002).

Insight into reproductive strategies of fungi have proven very important in population genetics and evolutionary studies of fungal pathogens. Sexual reproduction can play a key role in pathogen epidemiology, and determine the rate of evolution of important traits including fungicide resistance (Lopes et al., 2017; Meng et al., 2015). Although sexual structures are unknown for several species, population genetic analyses have provided evidence for the occurrence of sexual recombination in important plant pathogens and thereby provided essential knowledge regarding their epidemiology and evolutionary potential (Groenewald et al., 2006; McDonald and Linde, 2002; McDonald and McDermott, 1993)

Sexual reproduction has been studied in the heterothallic citrus pathogen *Phyllosticta citricarpa*, which causes the Citrus Black Spot (CBS) disease in sweet oranges and lemons. Ascospores of this fungus are responsible for infecting fruits and leaves and for long-distance dispersal of the pathogen (Kotzé, 1981). However, until recently, studies failed to induce sexual reproduction *in vitro* (Amorim et al., 2017; Baldassari et al., 2008; Wang et al., 2016). Successful production of ascospores under laboratory conditions was only possible after detailed analyses of the *MAT1* locus in *P. citricarpa*, which revealed the heterothallic nature of this species. Subsequently, PCR markers were developed to discriminate between *MAT1-1* and *MAT1-2* isolates (Amorim et al., 2017; Wang et al., 2016). Hereby, compatible isolates could be identified for *in vitro* crossings experiments and ascospore production (Tran et al., 2017).

Recent studies of the mating-type locus in *P. citricarpa* and a sister non-pathogenic species *P. capitalensis* (Wang et al., 2016) provided the foundation for a first comparative analysis of reproductive strategies in *Citrus*-associated *Phyllosticta* species. Although these two species are closely related, they have distinct mating strategies. While *P. citricarpa* is heterothallic, *P. capitalensis* is homothallic including both mating idiomorphs in the same isolate.

Genome sequences of closely related *Phyllosticta* species are a valuable resource for studying mating strategy evolution among species. To date there are eight *Phyllosticta* species known to be associated with *Citrus*, either as pathogens, e.g. *P. citricarpa, P. paracitricarpa, P. citrimaxima, P. citriasiana* and *P. citrichinaensis*, or as endophytes, e.g. *P. citribraziliensis, P. capitalensis* and *P. paracapitalensis* (Glienke et al., 2011; Guarnaccia et al., 2017; Wang et al., 2012; Wikee et al., 2013; Wulandari et al., 2009). The genomes of six of these species have been sequenced (Guarnaccia et al., 2019; Rodrigues et al., 2019; Wang et al., 2016). Comparative genome analyses combined with evidence for ascospore production in pure cultures (Glienke et al., 2011; Guarnaccia et al., 2017; Wang et al., 2012) provided evidence that *P. capitalensis, P. citrichinaensis* and *P. paracapitalensis* are homothallic, while *P. citricarpa, P. citriasiana, P. citribraziliensis* are heterothallic (Amorim et al., 2017; Guarnaccia et al., 2019; Wang et al., 2016).

This diversity in reproduction strategies, host-pathogen interactions, and lifestyles, makes the *Citrus*-associated *Phyllosticta* species a valuable model system to study the evolution of sexual mating strategies in fungal pathogens. Hereby, the main goal of this study was to elucidate the evolution of *Citrus*-associated *Phyllosticta* species’ mating strategies using a detailed characterization of the *MAT* loci. We present thorough analyses of gene order and sequence conservation along the mating-type locus for these *Phyllosticta* species, and make inferences about past shifts from homo- to heterothallism in the genus. We also discuss how sexual reproduction may have influenced the biology and evolution of *Citrus*-associated *Phyllosticta* species, and provide a foundation for further investigation of the evolutionary processes involved in these species’ diverse associations with hosts.

## 2 Materials and Methods

### 2.1 Genome data

Genome data used in the analyses were obtained from public assemblies available from GenBank (Wang et al., 2020; Wang et al., 2016) and the JGI MycoCosm portal (Guarnaccia et al., 2019). We also conducted *de novo* assemblies of both unpublished and already published data (Table 1). The published genomes of *P. capitalensis* LGMF01 and *P. citricarpa* LGMF06 are highly fragmented (Rodrigues et al., 2019), with the full mating-type locus not assembled. Therefore, we generated *de novo* genome assemblies for these strains using published Illumina mate-pair read data.

**Table 1.**
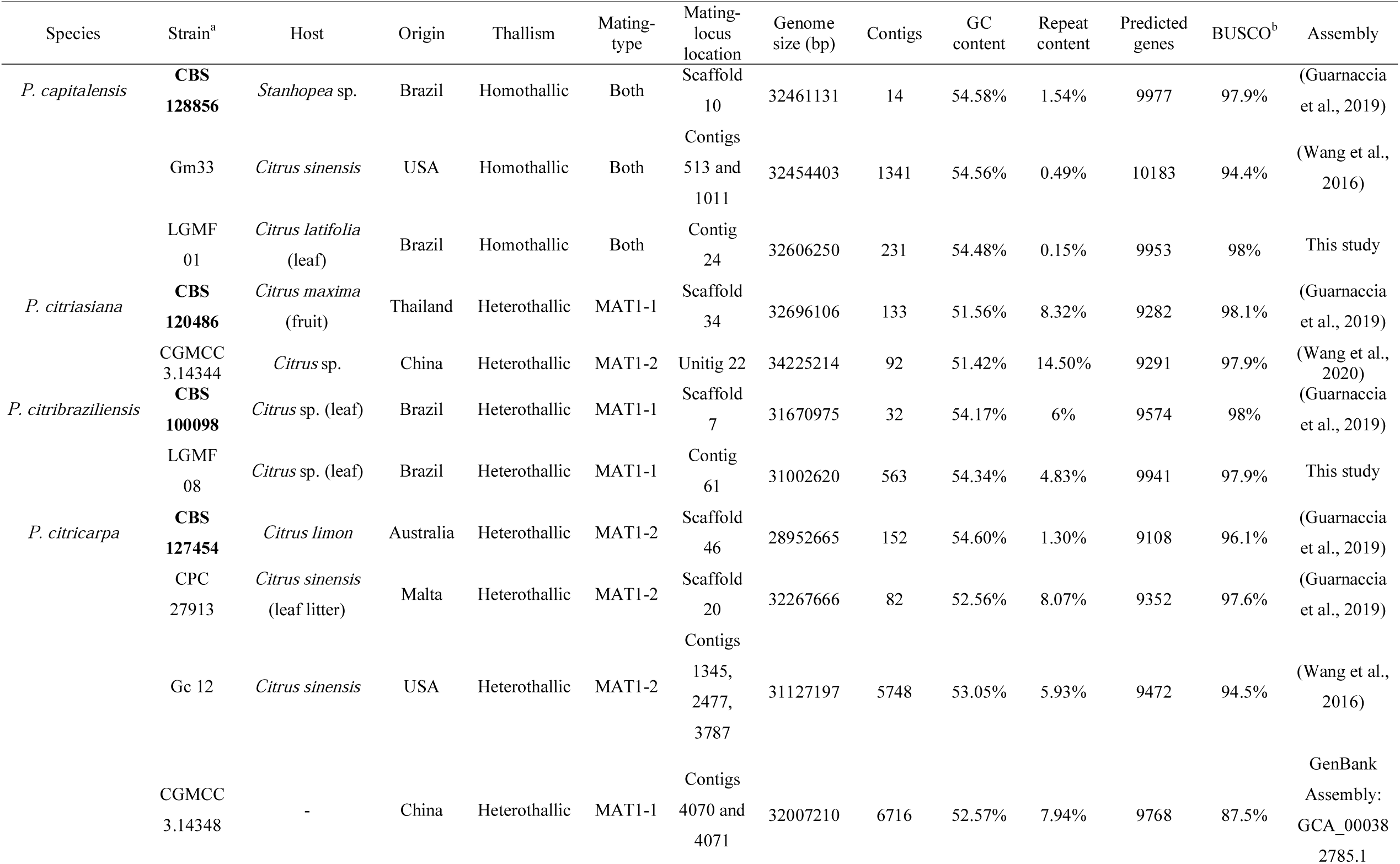
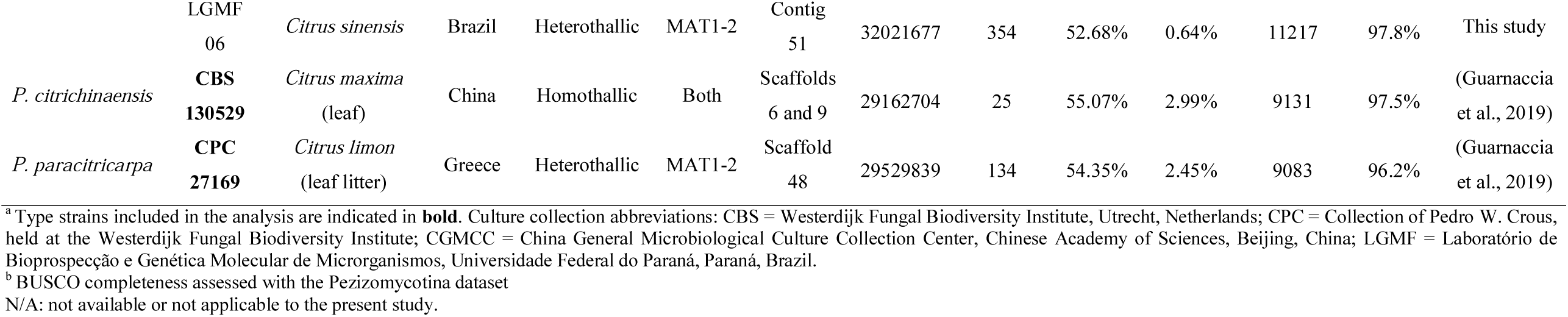
Genome and strain information of *Phyllosticta* spp. evaluated in this study

For *P. citribraziliensis* LGMF08, for which a public assembly was not available, genomic DNA was extracted from seven-day-old cultures grown in liquid potato-dextrose medium (PD) at 200 rpm and 28 °C, employing the CTAB method (Doyle and Dickson, 1987). Libraries were prepared with the Illumina Nextera DNA Sample Preparation Kit. Sequencing was performed using the Illumina HiSeq2500 platform, producing paired-end reads (250 bp), at the Laboratório Central de Tecnologias de Alto Desempenho em Ciências da Vida (LaCTAD), Campinas, Brazil.

For *P. capitalensis* LGMF01 and *P. citricarpa* LGMF06, libraries of the paired-end reads were processed with NxTrim (O’Connell et al., 2015) to remove Nextera adapters and generate mate-pair, paired-end, single-end and unknown libraries. The script “deinterleave_fastq.sh” (https://gist.github.com/nathanhaigh/3521724) was used to separate the reads from the mate-pair and paired-end libraries in “forward” and “reverse” files. For quality filtering, Trimmomatic v 0.38 (Bolger et al., 2014) was used to 1) trim bases at the start and end of the reads below a quality threshold of 25, 2) trim low quality segments using a 4 bp sliding window and a quality threshold of 15, and 3) discard reads shorter than 50 bp. For *P. citribraziliensis* LGMF08, the reads were filtered in Trimmomatic to remove Illumina adapters and trim for quality using the same parameters described before but discarding reads shorter than 90 bp. FastQC v 0.11.8 (https://www.bioinformatics.babraham.ac.uk/projects/fastqc/) was employed to check the quality of reads after processing in Trimmomatic. *De novo* genome assemblies were generated with SPAdes v 3.13 (Bankevich et al., 2012), using default parameters. The final assemblies were evaluated with QUAST v 4.6.3 (Gurevich et al., 2013). Library and assembly statistics for the new assemblies are summarized in Table S1 and assemblies are available at Zenodo (https://doi.org/10.5281/zenodo.3750350).

For a better understanding on how the genomes compare to each other, and to ensure quality and consistency for the downstream analyses, we assessed the repeat content and genome completeness levels of all genomes evaluated in this study. Repetitive regions were annotated with the REPET package v 2.5 (Flutre et al., 2011; Quesneville et al., 2005). First, we identified repetitive elements using TEdenovo with default parameters, which generates a library of classified and non-redundant consensus repeat sequences. Then, this library was used for the annotation of the genomes with TEannot. Genome completeness was assessed with BUSCO v 3 (Simão et al., 2015) using the Pezizomycotina lineage dataset (Waterhouse et al., 2018).

### 2.2 Gene annotation, <u>MAT</u> locus identification and characterization of mating-type and adjacent genes

Protein-coding genes were identified in *Phyllosticta* genome assemblies employing a similar strategy to the one used in the Fungal Genome Annotation pipeline (Haas et al., 2011). Briefly, GeneMark-ES v 4.46 (Ter-Hovhannisyan et al., 2008) was run in the self-training mode to provide an initial set of *ab initio* predictions. Then, to obtain a set of RNA-Seq-based predictions, RNA-Seq reads from the ex-type strains of the six *Phyllosticta* species analysed (Guarnaccia et al., 2019) were mapped to the genomes of the strains from the same species with HISAT2 v 2.1.0 (Kim et al., 2019). The mapping results were submitted to BRAKER1 (Hoff et al., 2016), which combines GeneMark-ET v 4.46 (Lomsadze et al., 2014) and AUGUSTUS v 3.3 (Stanke et al., 2008). In addition, genome-guided transcript reconstruction using Trinity v 2.8.4 (Grabherr et al., 2013; Haas et al., 2013) was performed. The generated transcripts were used by PASA v 2.4.1 to obtain a set of transcriptome-based gene models. Then, the EVidenceModeler software v 1.1.1 (Haas et al., 2008) was applied to generate the final weighted consensus set of gene structures, combining the *ab initio* and the transcript-based predictions. Gene predictions are available at Zenodo (https://doi.org/10.5281/zenodo.3750350).

The *MAT* locus was identified in each genome by a local tBLASTx search with BLAST v 2.2.26 applying the *Phyllosticta citricarpa MAT* idiomorphs and *APN2* genes (Amorim et al., 2017) as queries. The scaffold and contigs that contained sequences with high levels of similarity to the query sequences were extracted from the genomes, and the corresponding nucleotide and amino acid sequences of each gene were extracted based on the previously generated gene predictions. To confirm the identities of the predicted genes, the translated amino acid sequences were queried against the NCBI non-redundant (nr) database with BLASTp. After confirmation, the gene annotations were further refined by comparison with the *MAT* loci from *Phyllosticta citricarpa* and some *Botryosphaeriaceae* species (Nagel et al., 2018). The sequences and annotations are deposited in Zenodo (https://doi.org/10.5281/zenodo.3750350). Functional domains in the annotated genes were determined using the Conserved Domain Search Service (CD-Search) from NCBI (Lu et al., 2020), using an e-value of 0.01 as threshold.

### 2.3 Characterization and comparison of mating-type loci organization

Mating-type and flanking gene sequence similarities among *Phyllosticta* species were evaluated with a pairwise BLASTn analysis with a maximum e-value cut-off of 0.0001. The regions were then used for a synteny analysis in EasyFig v 2.2.3 (Sullivan et al., 2011). Intergenic regions were evaluated for the presence of partial *MAT1-1-1, MAT1-1-8, MAT1-2-1* and *MAT1-2-5* sequences. Finally, we assessed the presence of partial fragments of the mating-type locus in the rest of the genome.

### 2.4 Primer design, PCR and sequencing

In addition to the primers previously designed to amplify fragments from the *MAT1-1-1* gene of *Phyllosticta* species (Guarnaccia et al., 2019), new primers were designed to confirm the presence of the LTR-*Copia* element in the *P. citrichinaensis* mating-type locus.

Primers were designed based on genomic sequences from strain CBS 130529 and targeted the LTR-*Copia* element as well as the flanking gene *PIM*. The primers were tested for performance characteristics such as internal structures, hairpins, and self- and hetero-dimer formation using OligoIDTAAnalyzer v 3.1 (https://eu.idtdna.com/calc/analyzer/) (Owczarzy et al., 2008). Melting temperatures were calculated with the ThermoFisher Tm Calculator, to ensure primer compatibility. Primer sequences, annealing positions and amplicon sizes are summarized in Supplementary Table S2.

A DNA sample from *P. citrichinaensis* for PCR amplification was extracted with QIAGEN Genomic-tip 100/G (QIAGEN Benelux B.V., Venlo, Netherlands). PCR amplifications were performed with Phusion High Fidelity PCR Master Mix (Invitrogen, Carlsbad CA, USA). PCR conditions included an initial denaturation step of 98 °C for 30 s, 35 cycles of 98 °C for 10 s, annealing at 54 °C for 20 s, extension times of ∼30 s per kilobase of DNA, and a final extension step of 72 °C for 10 min. Amplicons were visualized on 0.8% agarose gel electrophoresis using 1× TAE buffer stained with SYBRSafe (Invitrogen, Carlsbad CA, USA), and compared to a GeneRuler 1 Kb Ladder (ThermoScientific, Waltham MA, USA) for evaluation of size, quality and concentration. For sequencing, amplicons were purified using illustra ExoProStar (GE Healthcare, Chicago IL, USA) and sequenced using the BigDye Terminator Cycle Sequencing Kit v 3.1 (Applied Biosystems, Foster City CA, USA). DNA sequences were obtained on an ABI 3130xl DNA sequencer (Applied Biosystems, Foster City CA, USA) and the electropherograms were examined and manually corrected when necessary using MEGA7 software (Kumar et al., 2016).

### 2.5 Phylogenetic analyses and ancestral state reconstruction

To provide a framework of species relationships for the synteny comparisons, we performed a multilocus analysis of the *ITS, TEF1, ACT* and *GAPDH* loci of *Citrus*-associated *Phyllosticta* species. Next, we investigated how the evolutionary history of mating-type and flanking genes relate to the evolutionary history of *Citrus*-associated species. To this end, we performed phylogenetic analyses of coding regions and amino acid sequences of *MAT1-1-1, MAT1-1-8, MAT1-2-1* and *MAT1-2-5*, as well as sequences from the flanking genes (*SAICARsyn, MCP, CIA30, CoxVIa, APN2, PIM and 40S S9*), using *Diplodia seriata* as outgroup. Finally, to provide a backbone for an ancestral state reconstruction of thallism, we performed a new multilocus analysis of *ITS, TEF1, ACT* and *GAPDH*, with eight additional representative species of the genus *Phyllosticta* and using *Diplodia corticola* and *Neofusicoccum parvum* as outgroups. Accession codes of all sequences used in the phylogenetic analyses are summarized in Table S3.

Alignments were produced in MAFFT v 7 (Katoh and Standley, 2013), and manually adjusted when necessary with the MEGA7 software (Kumar et al., 2016). Phylogenetic analyses were performed in the CIPRES Science Gateway portal v 3.3 (Miller et al., 2012) using two different approaches: Maximum Likelihood (ML) analysis with Garli v 2.0 (Zwickl, 2006) and Bayesian Inference (BI) analysis with MrBayes v 3.2.6 (Ronquist et al., 2012). Evolutionary models were tested using ModelTest-NG (Darriba et al., 2020) and the best-fit substitution model for each gene was selected based on the Akaike criterion (Supplementary Table S4).

For Maximum Likelihood analyses, evolution was simulated until likelihood scores converged. Nonparametric bootstrap analyses were performed with 1000 pseudoreplicates, in order to estimate the statistical support of the branches. Nodes with zero branch lengths were collapsed. Bootstrap trees were then compiled using SumTrees v 4.3.0 from the Dendropy v 4.4.0 package (Sukumaran and Holder, 2010). For Bayesian Inference, analyses were performed using two parallel runs and a sampling frequency set to every 10,000 generations; each of the two parallel runs had one cold and three heated chains, and were run until split frequencies were less or equal to 0.01. The first 25% of the generated trees were discarded as burn-in before calculating the 50% majority consensus trees for ML and posterior probability values for BI. The resulting trees from both analyses were outputted in FigTree v 1.4.4 (http://tree.bio.ed.ac.uk/software/figtree/) and their topologies were compared. We assessed possible incongruences and conflicts between clades with significant bootstrap/posterior probability support in both analyses, and compared their topologies to the ones presented in previously published phylogenies of *Phyllosticta* (Marin-Felix et al., 2019) and *Botryosphaeriales* (Phillips et al., 2018). Alignments and trees produced in this study are available in TreeBASE (study S25256, http://purl.org/phylo/treebase/phylows/study/TB2:S25256?x-access-code=7ecdbf532816b871453448fa454336e0&format=html).

The multilocus tree (*ITS*, T*EF1, ACT* and *GAPDH*) was used to map characters in the reconstruction of the ancestral thallic state in Mesquite v 3.61 (Maddison and Maddison, 2019). Character states were heterothallism (when only the *MAT1-1* or *MAT1-2* idiomorph was present in the genome), homothallism (when both mating-type idiomorphs were present in the genome, or when pure cultures were described to produce ascospores *in vitro*), or unknown. Besides the character states from *Phyllosticta* species, we also included the thallic states the outgroups, *Diplodia corticola* and *Neofusicoccum parvum* (Nagel et al., 2018).

## 3 Results

### 3.1 Genome assemblies and gene annotation

Our *de novo* assemblies for *P. capitalensis, P. citribraziliensis* and *P. citricarpa* resulted in genomes ranging from ∼31Mb for *P. citribraziliensis* to ∼32Mb for *P. capitalensis* and *P. citricarpa*, thereby being comparable to the other *Phyllosticta* genomes already available (Table 1). However, the assemblies revealed considerable differences in the extent of contig fragmentation: the new assemblies of *P. capitalensis* LGMF01 and *P. citricarpa* LGMF06 comprise 231 and 354 contigs, respectively, while the first published assemblies of these same strains included more than 11,000 contigs (Rodrigues et al., 2019).

An analysis of assembly completeness based on BUSCO demonstrates that our assemblies present genome completeness levels around 98%, comparable to the reference assemblies of the ex-type strains (Guarnaccia et al., 2019), regardless of being more fragmented (Table 1). This suggests that assembly fragmentation does not cause an increase in the number of missing protein coding genes. Instead, the unassembled fragments likely comprise repetitive sequences.

There is substantial variation in the repetitive content in *Phyllosticta* genomes. Some species present low TE content, such as *P. capitalensis* (0.14 to 1.54%), while other species present higher TE content, such as *P. citriasiana* strain CGM3.14344. The number of predicted genes varies between 9,083 (in *P. paracitricarpa* CPC 27169) and 11,217 genes (in *P. citricarpa LGMF06*) (Table 1).

### 3.2 Homothallism and heterothallism throughout the genus <u>Phyllosticta</u>

To enable a comparison of the mating-type locus structure in *Phyllosticta* species, we first assessed the presence of mating-type genes in the genomes. Genes with high similarity to mating-type genes of *P. citricarpa*, containing either the alpha-box domain (pfam04769) from the *MAT1-1-1* gene, or the high-mobility group (cd01389) domain from the *MAT1-2-1* gene, were found in all the analyzed *Phyllosticta* genomes.

The *MAT1-1-1* gene always included the alpha-box domain (Table 2) and co-occurred with the *MAT1-1-8* gene, together comprising the *MAT1-1* idiomorph. On the other hand, the *MAT1-2-1* gene always contained the HMG-box domain (Table 2), and co-occurred with the *MAT1-2-5* gene, comprising the *MAT1-2* idiomorph.

**Table 2.**
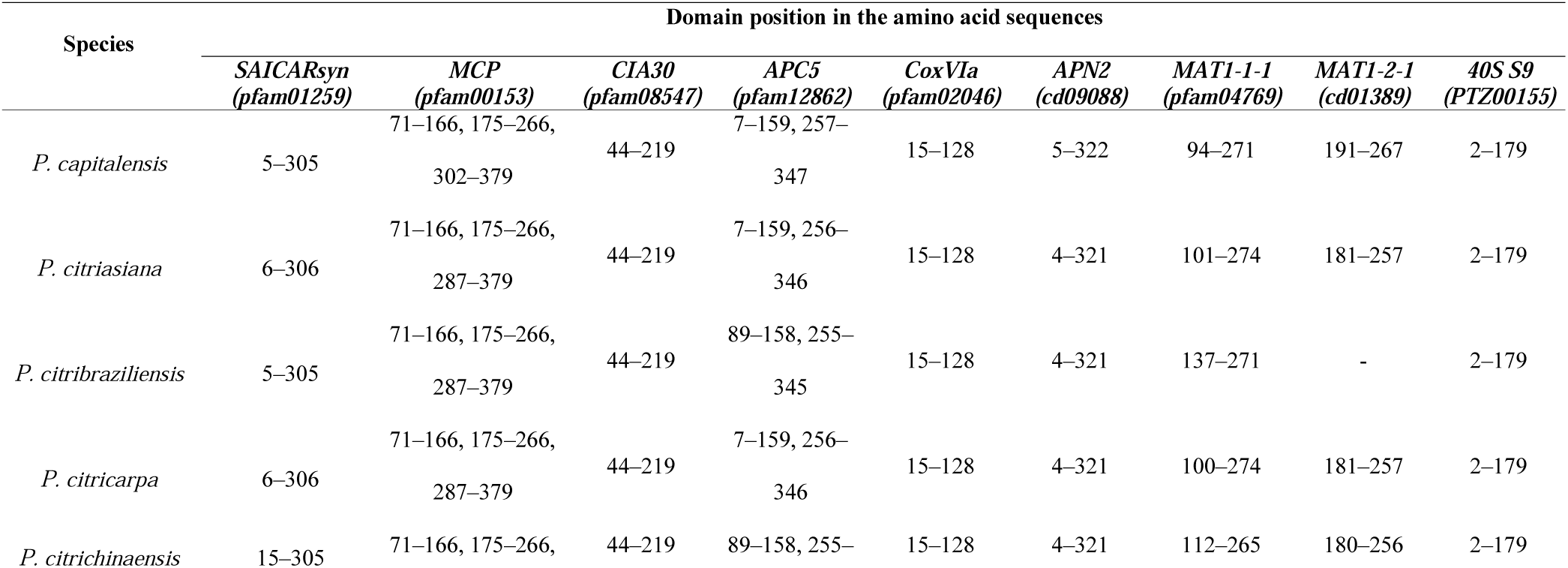
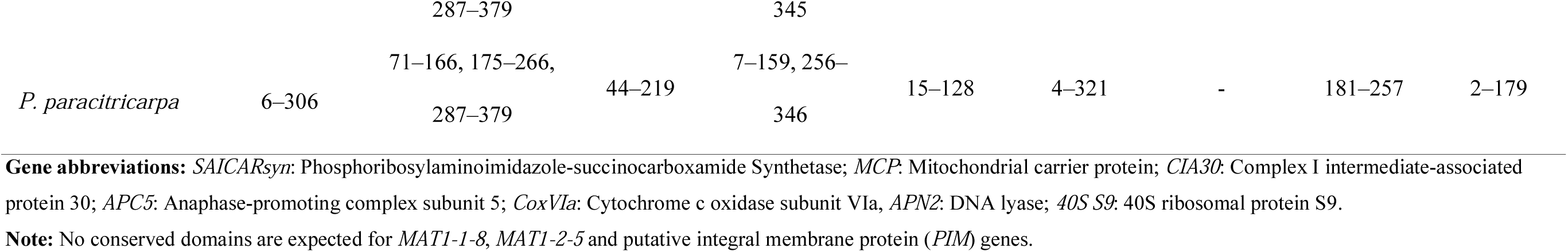
Conserved protein domains present in mating-type and flanking genes in *Phyllosticta* species

Based on the presence of mating-type idiomorphs, it is possible to classify *Phyllosticta* species as heterothallic (only *MAT1-1* or *MAT1-2* idiomorph in the genome) or homothallic (when both idiomorphs occur in the genome). Based on our analyses of mating-type idiomorphs we conclude that *P. capitalensis* and *P. citrichinaensis* are homothallic, and *P. citriasiana, P. citribraziliensis, P. citricarpa* and *P. paracitricarpa* are heterothallic (Table 1), All *P. citricarpa* strains (except strain CGMCC3.14348), *P. citriasiana* (CGMCC3.14344) and *P. paracitricarpa* (CBS 141357) encode the *MAT1-2* idiomorph, while the *P. citriasiana* strain CBS 120486 and the *P. citribraziliensis* strains encode the *MAT1-1* idiomorph (Table 1).

### 3.3 Mating-type locus conservation in <u>Citrus</u>-associated <u>Phyllosticta</u> species

To assess whether rearrangements in the mating-type locus are associated with transitions in mating strategies, we investigated the organization and synteny of the locus across the *Citrus*-associated *Phyllosticta* species. The analysis also included additional genes upstream and downstream of the mating-type locus. This enabled syntenic comparisons not only across the genus *Phyllosticta* (Phyllostictaceae), but also with the allied family Botryosphaeriaceae (Nagel et al., 2018).

The upstream region (∼12Kb in length) consists of a syntenic cluster of six genes (*SAICARsyn*-*APN2* cluster), which are all protein coding and are present in the following conserved order: Phosphoribosylaminoimidazole-succinocarboxamide synthetase (*SAICARsyn*), Mitochondrial carrier protein (*MCP*), Complex I intermediate-associated protein (*CIA30*), Anaphase-promoting complex subunit 5 (*APC5*), Cytochrome c oxidase subunit Via (*CoxVIa*) and DNA lyase (*APN2*). The downstream region (∼5Kb in length) comprises a gene coding a putative integral membrane protein (*PIM)* and the 40S ribosomal protein S9.

In relation to the mating-type genes, their order and orientation is also conserved in the *Citrus*-associated *Phyllosticta* species (Figure 1 and Figure S1). In the *MAT1-1* idiomorph of heterothallic species, *MAT1-1-8* was always followed by *MAT1-1-1*, in opposite orientations from each other. In the *MAT1-2* idiomorph, *MAT1-2-1* was always followed by *MAT1-2-5*, with opposite orientations from each other. Furthermore, when comparing both idiomorphs, *MAT1-1-1* and *MAT1-2-1* have opposite orientations. In the homothallic species *P. capitalensis*, which presents both idiomorphs in the same genomic location, the *MAT1-1* idiomorph was followed by the *MAT1-2* idiomorph, both between the *SAICARsyn*-*APN2* cluster and *PIM*.

**Figure 1.**
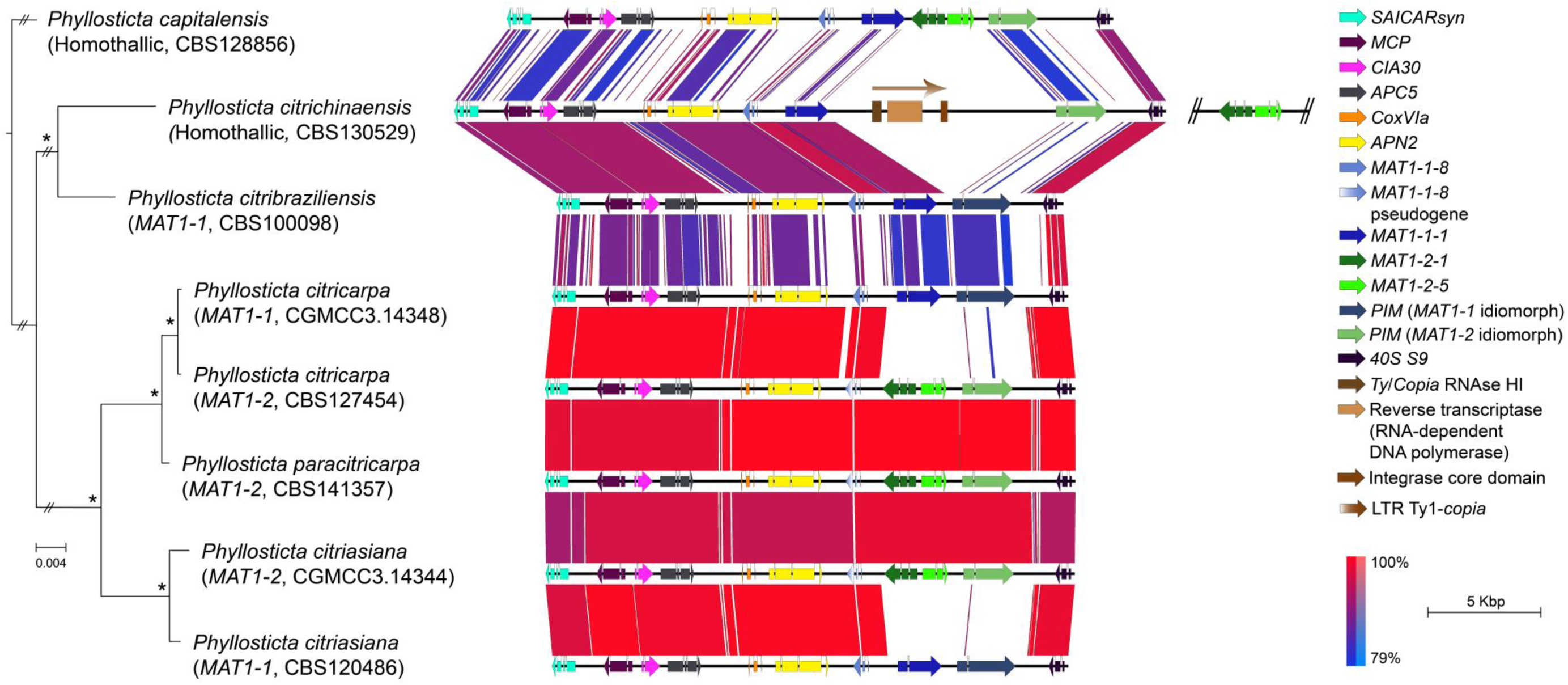
Pairwise comparison of mating-type and flaking genes between *Phyllosticta* species. The black horizontal lines represent nucleotide genomic sequences, and colour-coded arrows represent coding sequences in forward (right-oriented arrow) or reverse (left-oriented arrow) strand. Boxes between the genomic sequences represent the pairwise similarity based on BLASTn analyses, coded from red (most similar) to blue (least similar). Gene abbreviations: *SAICARsyn*: Phosphoribosylaminoimidazole-succinocarboxamide Synthetase; *MCP*: Mitochondrial carrier protein; *CIA30*: Complex I intermediate-associated protein 30; *APC5*: Anaphase-promoting complex subunit 5; *CoxVIa*: Cytochrome c oxidase subunit Via, *APN2*: DNA lyase; *PIM*: putative integral membrane protein; *40S S9*: 40S ribosomal protein S9. Phylogenetical relationships between *Phyllosticta* species, based in a multilocus dataset of the internal transcribed spacers and intervening 5.8S nrDNA (*ITS*), translation elongation factor 1-alpha (*TEF1*), actin (*ACT*), and glyceraldehyde-3-phosphate dehydrogenase (*GAPDH*) sequences are indicated by the cladogram on the left. ML bootstrap support values above 95% and Bayesian posterior probability values (PP) above 0.9 indicated with an asterisk (*) to the left of the nodes. Scale bar indicates the number of substitutions per nucleotide.

There are two different types of structure of the mating-type locus in the *Phyllosticta* species analysed (Figure 1 and Figure S1). In the first one, present in most of the species, the mating-type genes are localized between the upstream and downstream regions aforementioned. In this arrangement, the idiomorph-specific region has a size ranging from ∼5Kb in heterothallic species *P. citriasiana, P. citribraziliensis, P. citricarpa* and *P. paracitricarpa*, to ∼9Kb in the homothallic species *P. capitalensis* (Figure 1). The second type of structure comprises a different arrangement observed in the other homothallic species, *P. citrichinaensis*. The *SAICARsyn*-*APN2* cluster was found in scaffold 9 of the genome assembly, followed by the *MAT1-1* idiomorph, which was separated from *PIM* and 40S protein S9 (*40S S9*) by a ∼10Kb region rich in AT content (∼70%), with no predicted genes. Despite the absence of functional genes in this 10Kb region, degenerate domains of a reverse transcriptase (pfam07727), integrase (cl21549) and Ty/Copia RNAse HI (cd09272) were found by CDD-Search, implying the insertion of a transposable element into the mating-type locus. Moreover, the *MAT1-2* idiomorph was found in a different scaffold (scaffold 6) (Figure 1 and Figure S1). In this scaffold, only the *MAT1-2-1* and *MAT1-2-5* genes were present: 5’ and 3’ flanking sequences were not similar to any of the six genes from the syntenic cluster. Upstream to the *MAT1-2* idiomorph location, there is a ∼6Kb sequence identified as a TE fragment by REPET. However, the sequence is too degenerate to allow further characterization and recognition of conserved domains. Even though we could detect that the *MAT1-2* idiomorph was separated from the *MAT1-1* idiomorph and the *SAICARsyn*-*APN2* cluster, we were not able to determine if both idiomorphs are on the same or separate chromosomes.

### 3.4 <u>MAT1-2</u> strains present a truncated version of <u>MAT1-1-8</u> gene

Sequences showing similarity to the *MAT1-1-8* gene from *P. citricarpa* and *P. citriasiana* were observed in the intergenic region between *APN2* and *MAT1-2-1* genes in *P. citricarpa, P. citriasiana* and *P. paracitricarpa MAT1-2* strains (Figure S2). In the *MAT1-2* strains, *MAT1-1-8* sequences have a length of 1518 bp. Sequence comparisons with *MAT1-1-8* sequences of *MAT1-1* strains from the same species show 124 bp deletions in the 5’ end of the gene in *MAT1-2* strains. Moreover, there are 16 nucleotide substitutions along the sequence in *P. citricarpa* and *P. paracitricarpa*. These differences would lead to a truncated version of *MAT1-1-8* in *MAT1-2* strains, suggesting that it might be a pseudogene. Apart from these *MAT1-1-8* fragments, no fragments from *MAT1-1-1, MAT1-2-1* and *MAT1-2-5* were found in the opposite idiomorph, or in different genomic locations, for all the *Phyllosticta* species analysed.

### 3.5 <u>MAT</u> genes follow the evolutionary history of <u>Phyllosticta</u>, but <u>PIM</u> gene shows unexpected relationships

For all the genes, both Bayesian inference and maximum likelihood phylogenetical analyses provided similar topologies, and therefore we present single trees for each gene with combined results from both approaches.

For the six genes in the flanking syntenic cluster (*SAICARsyn, MCP, CIA30, APC5, CoxVIa* and *APN2*) (Figures S3 to S8), and for the mating-type genes (*MAT1-1-1, MAT1-1-8, MAT1-2-1*) (Figure 2, Figures S9 to S11), both for nucleotide and amino acid sequences, the trees are highly similar to the recognized species tree inferred from the multilocus phylogeny using ITS, *TEF1, ACT* and *GAPDH*, as well as previously published phylogeny (Marin-Felix et al., 2019) and phylogenome (Guarnaccia et al., 2019) of the genus *Phyllosticta*. The exceptions are *MAT1-2-*5 (Figure S12), for which it was not possible to observe resolution at species level for *P. citricarpa, P. citriasiana* and *P. paracitricarpa*, and *40S S9* (Figure S13), which is highly conserved across the genus *Phyllosticta*.

**Figure 2.**
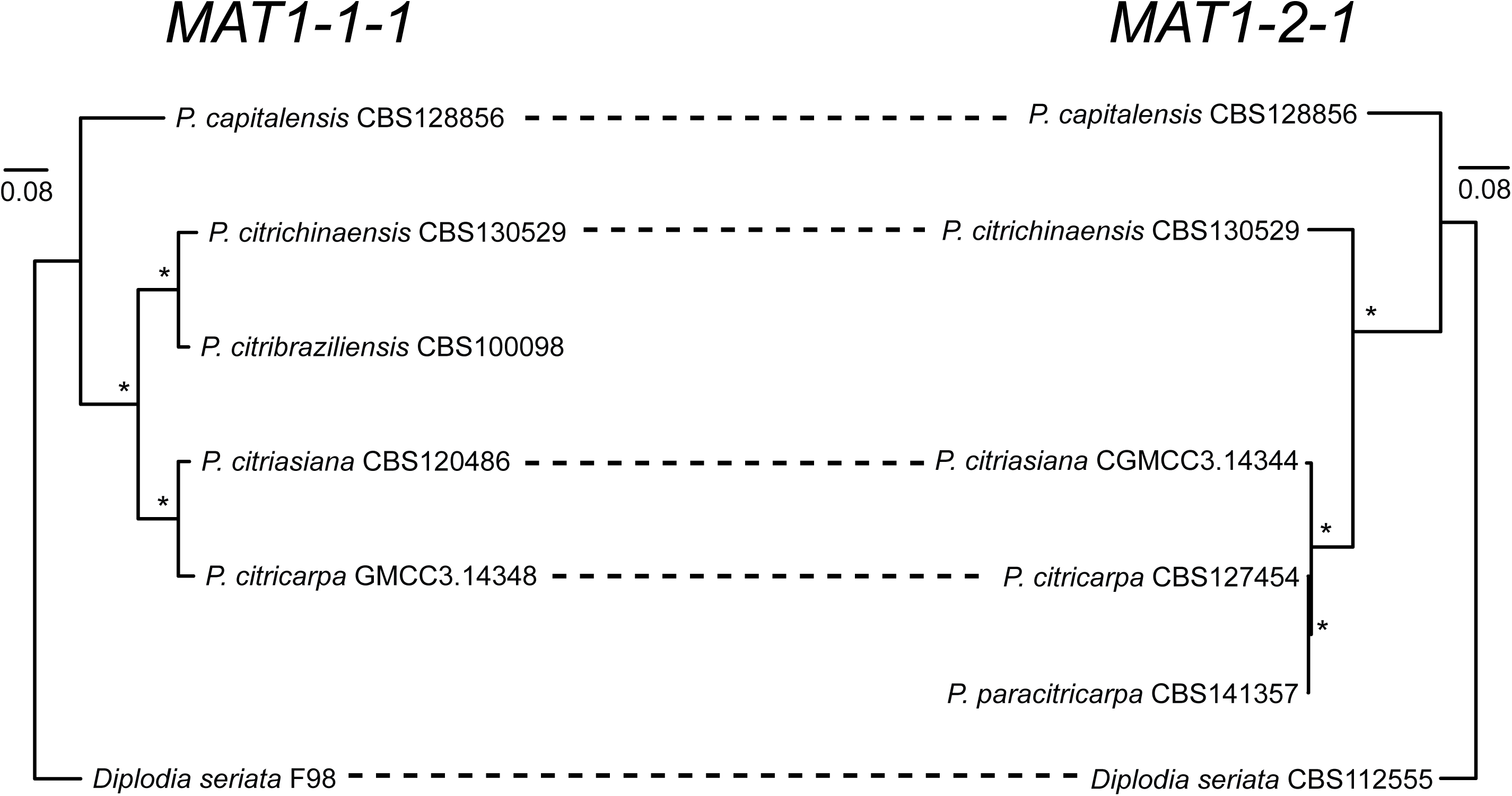
Bayesian Inference tree of *MAT1-1-1* and *MAT1-2-1* nucleotide sequences of *Phyllosticta* species. ML bootstrap support values above 95% and Bayesian posterior probability values (PP) above 0.9 indicated with an asterisk (*) to the left of the nodes. Both trees are rooted to *Diplodia seriata*. Scale bar indicates the number of substitutions per nucleotide.

However, the *PIM* gene was unexpectedly incongruent with the species tree (Figure 3, Figure S14). The *PIM* sequences of *P. citricarpa* strain CGM3.14348 and *P. citriasiana* strain CBS 120486 (both *MAT1-1* strains) group together in a separate clade, instead of grouping with the *PIM* sequences from the other isolates of their respective species. Furthermore, the species tree inferred from *PIM* sequences shows a different relationship for *P. citribraziliensis* and *P. citrichinaensis*. The multilocus species tree (ITS+*TEF1*+*ACT*+*GAPDH*) (Figure 1) shows *P. citribraziliensis* being most closely related to *P. citrichinaensis*, and both are in a separate clade from *P. citriasiana, P. citricarpa* and *P. paracitricarpa*. However, in the tree inferred from *PIM* sequences, *P. citribraziliensis* is not most closely related to *P. citrichinaensis* as expected, but groups in the same clade as *P. citriasiana* and *P. citricarpa MAT1-1* strains. This suggests that there are two different versions of the *PIM* gene, and that these versions show some level of idiomorph specificity: for heterothallic species, *PIM* genes group according to the idiomorph of the strain. As for the homothallic species, for *P. capitalensis* the *PIM* gene is closely related to the *PIM* from *MAT1-2* strains. This could be related to the mating-type locus organization in *P. capitalensis* where *PIM* is located right next to the *MAT1-2* idiomorph genes. Interestingly, the *P. citrichinaensis PIM* also groups with the *PIM* from *MAT1-2* strains, even though the *MAT1-2* idiomorph is in another genomic location.

**Figure 3.**
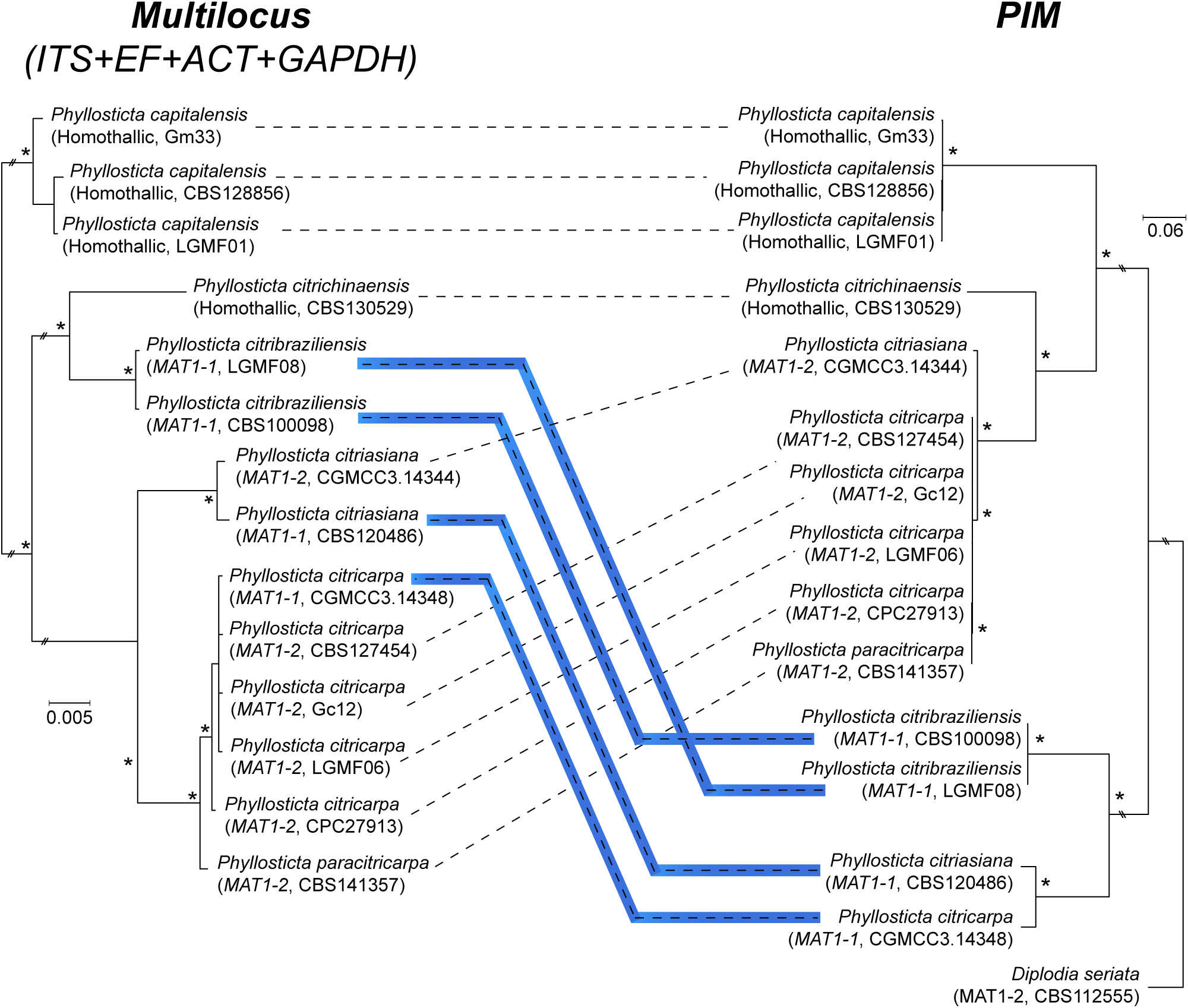
Bayesian Inference tree of a multilocus dataset (*ITS*+T*EF1*+*ACT*+*GAPDH*) and *PIM* nucleotide sequences of *Phyllosticta* species. ML bootstrap support values above 95% and Bayesian posterior probability values (PP) above 0.9 are indicated with an asterisk (*) to the left of the nodes. The *PIM* tree is rooted to *Diplodia seriata*. Scale bar indicates the number of substitutions per nucleotide. Conflicts between the species tree based on the multilocus dataset and the tree based on *PIM* are highlighted in blue.

### 3.6 The ancestral mating type locus of the genus <u>Phyllosticta</u> was homothallic

To assess the direction of transition between homothallic and heterothallic forms we performed an ancestral state reconstruction analysis. Hereby we find evidence for homothallism being the ancestral form in the genus *Phyllosticta* (Figure 4). Homothallism is present in several species from different clades, while heterothallism is restricted to a clade comprised mostly of *Citrus*-associated species.

**Figure 4.**
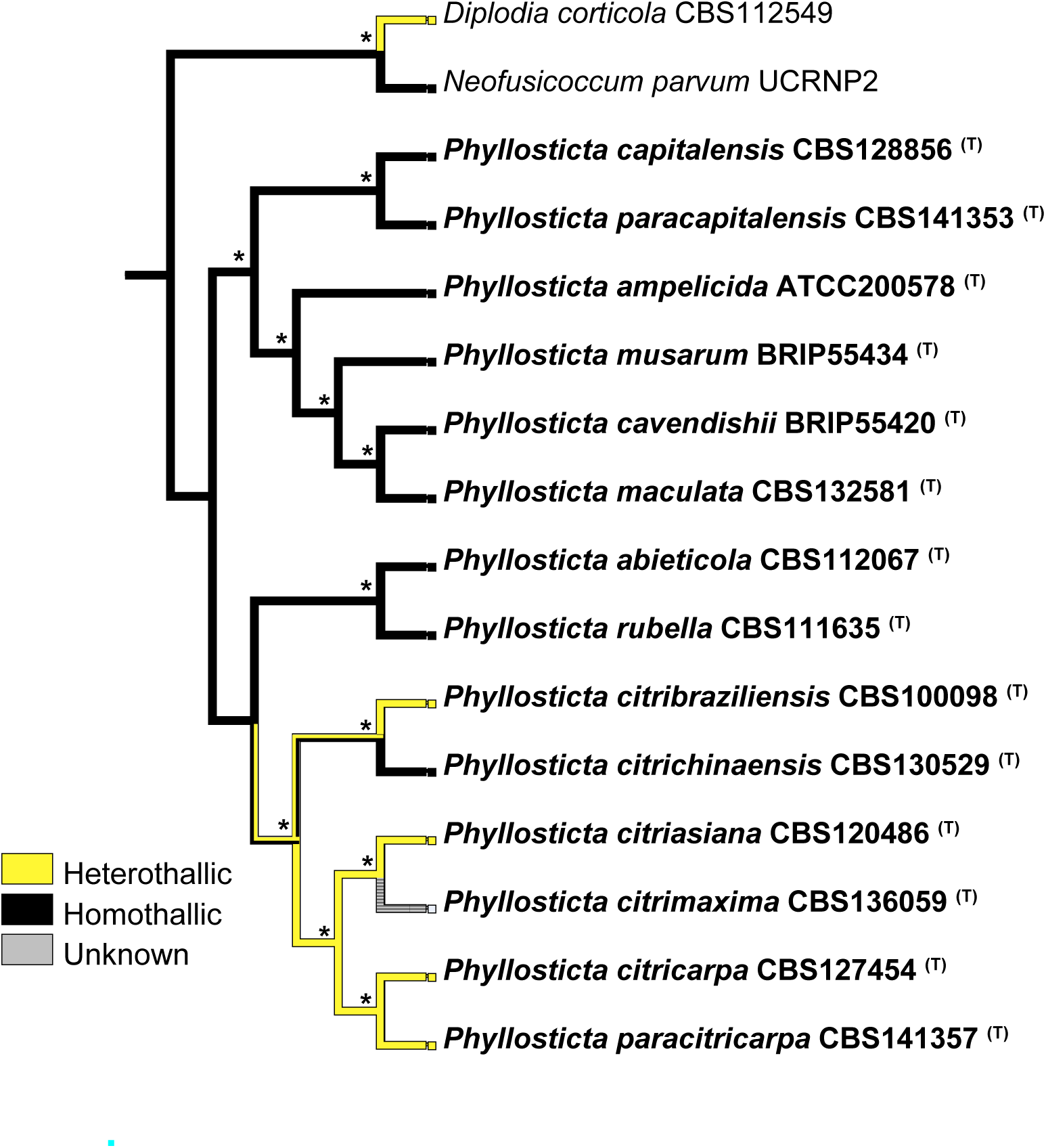
Ancestral state reconstruction of thallism states in *Phyllosticta* species. Characters were mapped onto a multilocus phylogenetic tree based on *ITS, TEF1, GAPDH* and *ACT* sequences. Thallism states are colour-coded according to the legend next to the tree. ML bootstrap support values above 95% and Bayesian posterior probability values (PP) above 0.9 are indicated with an asterisk (*) to the left of the nodes. The tree is rooted to *Diplodia corticola* and *Neofusicoccum parvum*. Scale bar indicates the number of substitutions per nucleotide.

Moreover, the reconstruction suggests that homothallism of *P. capitalensis* and *P. citrichinaensis* does not originate from the same evolutionary event. This scenario is also supported by the fact that they differ in the mating-type locus organization: in *P. capitalensis*, both idiomorphs are present in the same locus, whereas in *P. citrichinaensis* only the *MAT1-1* is present at the main mating-type locus, and *MAT1-2* occurs in another genomic region.

### 3.7 Mating-type genes present a different rate of sequence evolution

To investigate if the mating-type genes present a different rate of evolution from genes which are not associated with reproduction processes, we compared the degree of conservation, both at nucleotide and amino acid level, for the mating-type and flanking genes. Nucleotide and amino acid sequences of the *MAT1-1-1, MAT1-1-8, MAT1-2-1* and *MAT1-2-5* genes are less conserved than the sequences from the flanking genes (Table 3). The similarity level for mating-type genes ranges from 64.89% to 75.97% for nucleotide and from 53.19% to 61.94% for amino acid sequences. In contrast, for the flanking genes, the similarity level is higher, ranging from 69.30% to 93.21% for nucleotide and from 70.22% to 98.95% for amino acid sequences. Interestingly, despite being a flanking gene, the similarity level at the *PIM* gene is lower than the mating-type genes: 53.07% for nucleotide and 43.45% for amino acid sequences. This low similarity level is explained by the differences presented by the *PIM* sequences from *P. citribraziliensis, P. citriasiana* and *P. citricarpa MAT1-1* strains, when compared to the rest of the *PIM* sequences. This reinforces the suggestion of two versions of the *PIM* gene being present, as seen by the phylogenetic analysis (Figure 3).

**Table 3.** Nucleotide and amino acid conservation of mating-type and flanking genes in *Citrus*-associated *Phyllosticta* species

| Gene | Nucleotide |  | Amino acid |  |
| --- | --- | --- | --- | --- |
|  | *Conserved<sup>a</sup> / total sites (bp)* | *Sequence similarity (%)<sup>b</sup>* | *Conserved<sup>a</sup> / total sites (aa)* | *Sequence similarity (%)<sup>b</sup>* |
| *SAICARsyn* | 693/890 | 77.87% | 233/304 | 76.64% |
| *MCP* | 891/1149 | 77.55% | 342/382 | 89.53% |
| *CIA30* | 573/698 | 82.09% | 199/235 | 84.68% |
| *APC5* | 1621/2339 | 69.30% | 547/779 | 70.22% |
| *CoxVIa* | 346/392 | 88.27% | 108/130 | 83.08% |
| *APN2* | 1386/1914 | 72.41% | 479/642 | 74.61% |
| *MAT1-1-1* | 1169/1727 | 67.69% | 328/589 | 55.69% |
| *MAT1-1-8* | 621/957 | 64.89% | 175/329 | 53.19% |
| *MAT1-2-1* | 949/1296 | 73.23% | 275/444 | 61.94% |
| *MAT1-2-5* | 708/932 | 75.97% | 191/312 | 61.22% |
| *PIM* | 1003/1890 | 53.07% | 292/672 | 43.45% |
| *40S S9* | 535/574 | 93.21% | 189/191 | 98.95% |

<sup>a</sup> Identical sites in the alignment of *Citrus*-associated *Phyllosticta* species, excluding gaps
<sup>b</sup> Sequence similarity based in the proportion of identical sites in relation to the total number of sites in the alignment

On the other hand, despite the high variation at sequence level, we observed a high degree of conservation in terms of gene structure, for both the mating-type and adjacent genes. This conservation is observed at the size, number and position of exons and introns. One of the few exceptions would be the *MAT1-1-8* gene in *P. citricarpa* and *P. citriasiana MAT1-1* strains, in which the first intron is larger (∼267 bp) than the introns from its homologues in *P. capitalensis, P. citribraziliensis* and *P. citrichinaensis* (∼48 bp).

## Discussion

In our study we performed a detailed and comparative characterization of the mating-type locus in *Citrus*-associated *Phyllosticta* species. Moreover, we provide important resources for future comparative and evolutionary studies in this group of fungal species, in the form of new robust whole genome assemblies and gene annotations. Here, our comparative analyses demonstrate examples on how specific rearrangements in the overall conserved mating-type locus of *Phyllosticta* species may lead to important biological changes, such as the shift from homothallism to heterothallism.

Following the usual BLAST-based approach employed as the starting point in studies on mating-type loci in fungal species, we could identify the presence of *MAT* idiomorphs in all genomes analysed. Besides the essential *MAT1-1-1* and *MAT1-2-1* genes, which per se are the characteristic features of the canonical *MAT* loci in Ascomycetes, we demonstrate the presence of additional genes in the loci of both idiomorphs. A similar situation was also found in genera of the *Botryosphaeriaceae*, which is a family sister to *Phyllostictaceae* (Botryosphaeriales, Dothideomycetes). In *Botryosphaeriaceae* species such as *D. sapinea* (Bihon et al., 2014), *Neofusicoccum* spp. (Lopes et al., 2017), *Botryosphaeria dothidea, Macrophomina phaseolina* and *Lasiodiplodia* spp. (Nagel et al., 2018), *MAT1-1-8* and *MAT1-2-5* co-occur with *MAT1-1-1* and *MAT1-2-1*, respectively.

*MAT1-1-8* was previously referred to as *MAT1-1-4* in *P. citricarpa* (Amorim et al., 2017) and *oml1* (Wang et al., 2016) in *P. citricarpa* and *P. capitalensis. MAT1-1-4* was originally described in *Pyrenopeziza brassicae* as a mating-type gene with homology to metallothionein proteins (Wilken et al., 2017). These proteins are usually associated with the regulation of cellular growth and homeostatic control of heavy metal levels, suggesting that *MAT1-1-4* expression is induced by the senescence of host leaves, a situation in which there is an accumulation of heavy metal ions. This would be particularly important for the reproduction of *Py. brassicae*, which produces sexual structures in senescing tissue of the host (Wilken et al., 2017). *MAT1-1-8* was previously assigned as *MAT1-1-4* due to 31% of sequence identity to the *MAT1-1-4* gene described in *Diplodia sapinea* (Bihon et al., 2014). However, a recent revision of the mating-type genes nomenclature suggests that the gene from *D. sapinea* does not meet the required criteria to be classified as *MAT1-1-4* (Wilken et al., 2017). Firstly, there is the low sequence similarity to the original *MAT1-1-4* described in *Py. brassicae* (below 20%) and secondly, there is no information regarding the existence of a possible functional metallothionein domain in the *D. sapinea* gene. Given this, the authors propose that both are likely different genes, and that *MAT1-1-4* from *D. sapinea* be renamed as *MAT1-1-8* instead, allowing for future investigation and determination of the *MAT1-1-8* gene product and its biological function (Wilken et al., 2017). Thus, we propose here to adapt the *Phyllosticta* mating-type nomenclature to accommodate this change.

Likewise, *MAT1-2-5* was previously referred to as *MAT1-2-9* in *P. citricarpa* and *P. capitalensis* (Wang et al., 2016), due to the use of *MAT1-2-5* as a name for an idiomorph-specific *COX13* version in *Coccidioides* species (Mandel et al., 2007). Nomenclature of this gene was also revised and corrected, with the proposition of retaining *MAT1-2-5* for *P. citricarpa* and. *D. sapinea*, and referring to *Coccidioides* genes as alternative versions of *COX13* instead (Wilken et al., 2017), which we follow accordingly. As is the case for *MAT1-1-8*, further studies on the *MAT1-2-5* biological function and involvement in sexual reproduction are still required.

Regarding the overall mating-type locus organization, the presence of the synteny cluster *SAICARsyn*-*APN2* upstream to the mating-type idiomorphs, in this orientation, is also observed in different *Botryosphaeriaceae* species, such as *Lasiodiplodia pseudotheobromae, L. gonubiensis, Botryosphaeria dothidea* and *Macrophomina phaseolina* (Nagel et al., 2018). This high degree of similarity spanning from *Phyllosticta* across the whole order *Botryosphaeriales* is consistent with the frequent descriptions of positional conservation for the mating-type locus, commonly associated to genes such as *APN2* (Wilken et al., 2017).

However, despite the high level of overall organization for the mating-type locus, a remarkable rearrangement was observed for *P. citrichinaensis*, as one of the idiomorphs occurs in a separate genomic location. Comparable to this different kind of arrangement, previous studies have documented the presence of mating-type genes separate from the main mating-type locus, e.g. in *Aspergillus nidulans* (Galagan et al., 2005), *Eutiarosporella darliae* (Thynne et al., 2017), *Neurospora sublineolata* (Gioti et al., 2012), *Neofusicoccum australe* and *N. luteum* (Nagel et al., 2018). Moreover, *in Cercospora beticola*, apart from the main complete mating-type locus, fragments of *MAT1-1-1* and *MAT1-2-1* are dispersed throughout the genome (Bolton et al., 2014), something we did not observe for the *Citrus*-associated *Phyllosticta* species. Interestingly, the mechanisms involved in the remodelling of the mating-type locus in the aforementioned species differ. For example, in the heterothallic species *C. beticola* the *MAT* fragments do not contain introns, which implies that they likely originated from RNA transcripts that integrated themselves in the genome (Bolton et al., 2014). On the other hand, in the homothallic species *E. darliae* the *MAT1-1-1* gene, which is separated from the main *MAT* locus, still retains the intron sequences, as well as flanking fragments of non-coding DNA. In this case, it has been proposed that a DNA breakage or transposition event is responsible for the excision of *MAT1-1-1* from an ancestor heterothallic strain, which was later integrated in the *E. darliae* genome (Thynne et al., 2017). For *N. sublineolata*, the presence of the repetitive element *nsubGypsy* inside the mating-type locus suggests that the rearrangement of this locus happened through a TE-mediated translocation (Gioti et al., 2012). However, despite the several possible mechanistic differences observed in these species, one common outcome of the mating-type locus remodelling is a shift in thallism state, usually from heterothallism to homothallism. Studies in Ascomycetes show that the change from heterothallism is a frequent event (Nagel et al., 2018; Thynne et al., 2017), and may occur multiple times within one genus (Gioti et al., 2012). Moreover, it has been proposed that shifts from heterothallism to homothallism could have been a driver of speciation among pathogenic *Botryosphaeriaceae* species (Thynne et al., 2017).

Our results suggest that what happened in the genus *Phyllosticta* was the opposite: a shift from homothallism to heterothallism. Due to the basal placement of *P. capitalensis* in the phylogenetic analyses (Figure 4; also see Wikee et al., 2013), we hypothesize that the mating-type locus organization for the homothallic ancestor of the *Phyllosticta* genus would likely resemble what is observed for *P. capitalensis*, with both *MAT* idiomorphs occurring in the same locus. From this homothallic ancestor, we propose two models to explain the mating-type locus organization for the homothallic species *P. citrichinaensis* and for the heterothallic species *P. citriasiana, P. citribraziliensis, P. citricarpa* and *P. paracitricarpa* (Figure 5).

**Figure 5.**
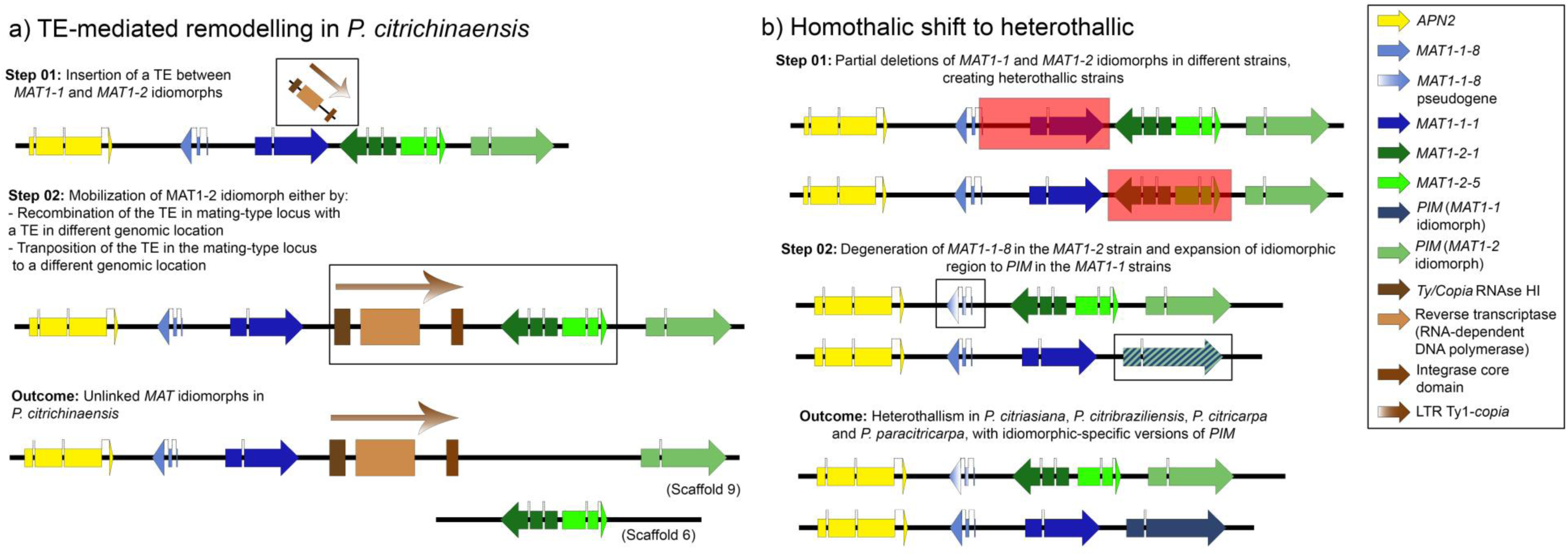
Mating-type locus rearrangement in Citrus-associated Phyllosticta species. (A) Schematic representation of TE-mediated remodeling in *P. citrichinaensis*. 1) Insertion of a TE between *MAT1-1* and *MAT1-2* idiomorphs, in the ancestor with a *P. capitalensis*-like *MAT* locus. 2) Mobilization of the *MAT1-2* idiomorph could have happen either by: 2a) the recombination of the TE in the mating-type locus with a TE in a different genomic location, 2b) the transposition of the TE in the mating-type locus to a different genomic location, moving the flanking sequence containing the *MAT1-2* idiomorph. 3) This rearrangement results in unlinked *MAT* idiomorphs in *P. citrichinaensis*. (B) Schematic representation of the homothallic to heterothallic shift in *P. citriasiana, P. citribraziliensis, P. citricarpa* and *P. paracitricarpa*. 1) Partial deletions of *MAT1-1* and *MAT1-2* idiomorphs in separated strains from the ancestor with a *P. capitalensis*-like *MAT* locus, creating heterothallic individuals. 2) Degeneration of the remaining *MAT1-1-8* gene in the *MAT1-2* strains, and expansion of the idiomorphic region until the *PIM* gene in *MAT1-1* strains. 3) These changes result in heterothallism in *P. citriasiana, P. citribraziliensis, P. citricarpa* and *P. paracitricarpa*, and idiomorphic-specific versions of *PIM*. The black horizontal lines represent nucleotide genomic sequences, and colour-coded arrows represent coding sequences in forward (right-oriented arrow) or reverse (left-oriented arrow) strand. Gene abbreviations: *APN2*: DNA lyase; *PIM*: putative integral membrane protein.

In *P. citrichinaensis*, we hypothesize a model in which the mating-type locus remodelling was mediated by TEs (Figure 5a). First, an insertion of a transposable element between *MAT1-1* and *MAT1-2* idiomorphs occurred. Then, this event was followed either by a recombination event between the TE inserted in the mating-type locus and another TE in a different genome location (e.g. the degenerate TE found ∼6Kb upstream the *MAT1-2* idiomorph location), or solely by the mobilization of the TE. The consequence of this translocation would be the mating-type locus organization that is observed in *P. citrichinaensis*, in which *MAT1-2* is separated from the rest of the mating-type locus. This model also explains the phylogenetic placement of the *P. citrichinaensis PIM* together with the *PIM* genes of *MAT1-2* strains (Figure 3): even though there is no physical association between the *PIM* gene and *MAT1-2* genes in this species anymore, the *MAT1-2* idiomorph was closely located to the *PIM* gene in the *P. citrichinaensis* ancestor (Figure 5a). The involvement of TEs in the remodelling of the mating-type locus has already been proposed in other species such as *Neurospora sublineolata* (Gioti et al., 2012), *Chrysoporthe austroafricana* (Kanzi et al., 2019) and *Sclerotinia trifoliorum* (Xu et al., 2016), either through translocations, gene loss and unidirectional mating-type switching. Moreover, TEs are known to impact the genome evolution of several fungal species by promoting chromosomal rearrangements (Castanera et al., 2016). Even though the TE content in *Phyllosticta* species seems to be low, it is possible that some of these TEs may be involved in important rearrangements, such as the one we describe here for the mating-type locus. In this sense, further characterization on the TE dynamics in the *Phyllosticta* species may provide interesting insights.

For the heterothallic species *P. citriasiana, P. citribraziliensis, P. citricarpa* and *P. paracitricarpa*, we propose a model based on gene loss (Figure 5b). In different individuals from a homothallic ancestor, partial deletions of either the *MAT1-1* or *MAT1-2* idiomorphs occurred, creating heterothallic strains incapable of selfing. In the *MAT1-1* strains, the whole *MAT1-2* idiomorph was lost whereas, in the *MAT1-2* strains, only the *MAT1-1-1* gene was completely lost, as a truncated version of *MAT1-1-8* still exists. This presence of partial *MAT1-1-8* in *MAT1-2* strains is not unusual, and is already reported in literature for some other *Botryosphaeriales* species, such as *Diplodia pinea* (Bihon et al., 2014), *Lasiodiplodia theobromae, L. gonubiensis, Macrophomina phaseolina, Neofusicoccum australe* and *N. luteum* (Nagel et al., 2018). Moreover, it has been proposed that these partial gene fragments in the mating-type locus of heterothallic strains can arise either from heterothallic ancestors that underwent unequal crossing over/recombination events, or from homothallic ancestors that experienced two independent deletion events (Nagel et al., 2018), such as what we hypothesize for the genus *Phyllosticta*. In this scenario, it is possible to expect some level of homologous recombination due to the considerable sequence similarity between the *MAT1-1-8* gene and the partial fragments in the *MAT1-2* idiomorph in *Phyllosticta* species. This recombination would reduce the beginning of the effective idiomorphic region (Nagel et al., 2018) to just part of the *MAT1-1-8* gene.

Nevertheless, an opposite effect is observed for the *PIM* gene which includes two idiomorph-specific versions (Figures 1 and 3). This diversity suggests that *PIM* is being incorporated into the idiomorphic region not only in *P. citricarpa* (Wang et al., 2016), but also in the other heterothallic species. The expansion of the idiomorphic region has already been observed for other fungal species, even including genes that are not involved in the mating-type determination (Branco et al., 2017; Wang et al., 2016). To some extent, the expansion of the recombination suppression region observed in the heterothallic *Phyllosticta* species is consistent with proposed models of evolution of *MAT* loci towards sex chromosomes, which occurs through successive increases of the recombination suppression regions (evolutionary strata) (Branco et al., 2017).

The incorporation of the *PIM* gene to the idiomorphic region is also evidenced by the low degree of sequence conservation, comparable to what is observed for the mating-type genes. Mating-type genes usually evolve at faster rates than other sequences in the genome (Turgeon, 1998), contrasting with the conservation observed at gene structure and genomic levels for the mating-type locus. This diversity in evolutionary rates reinforces the importance of evaluating the genome evolution of the *Citrus*-associated *Phyllosticta* species across different levels.

Our characterization of the mating-type locus in the *Citrus*-associated *Phyllosticta* species also provides foundations for further functional studies on the sexual reproduction processes in these species. As mentioned before, *MAT1-1-8* and *MAT1-2-5* genes still lack information regarding their biological products and functions (Wilken et al., 2017). Moreover, little is known about expression of mating-type genes in *Phyllosticta* species, especially in the context of the interaction with the hosts. In this sense, functional approaches focusing on gene expression and characterization (Kim et al., 2015; Wilson et al., 2020) are required to determine the exact contribution of *MAT1-1-1, MAT1-1-8, MAT1-2-1* and *MAT1-2-5* to sexual reproduction in *Phyllosticta*.

The investigation of the sexual reproduction in *Phyllosticta* species also provides valuable insights in the context of the Citrus Black Spot and the Citrus Tan Spot diseases, by allowing an evaluation of ascospore pathogenicity and its role in the disease cycle. Regarding *P. citricarpa*, ascospore production and pathogenicity were described on leaves of Troyer seedlings and fruits of Murcott tangor (Tran et al., 2018, 2017). Moreover, *P. citricarpa* isolates with different genotypes vary in their aggressiveness, implying that the genetic diversity generated by sexual reproduction has epidemiological significance and should be taken into consideration (Tran et al., 2018). Regarding the other pathogenic species, further investigation on ascospore pathogenicity and the influence of genetic diversity in disease development are still required. For *P. citriasiana*, pathogenicity assays were not conducted since its description (Wulandari et al., 2009), and for *P. paracitricarpa*, only conidia were evaluated until now (Guarnaccia et al., 2017). As both species are heterothallic, similar strategies to the ones employed in *P. citricarpa* (mating-type PCR and crossings with opposite isolates) (Amorim et al., 2017; Guarnaccia et al., 2019; Tran et al., 2017; Wang et al., 2016) may allow the *in vitro* production of ascospores and their evaluation in future. As recombination increases genetic diversity and can generate new combinations of virulence alleles, disease management practices that prevent the occurrence of sexual reproduction could assist in the control of *P. citriasiana, P. citricarpa* and *P. paracitricarpa*, similar to other recombining pathogens (McDonald and Linde, 2002; McDonald and Mundt, 2016). The successful implementation of such practices relies on the knowledge about the mating-type distribution in different populations, reinforcing the need of a broader sampling of isolates from these *Phyllosticta* species (Guarnaccia et al., 2019).

Finally, interesting insights on the link between speciation and shifts between homothallism and heterothallism may become available from further investigations on mating-type loci from other species of *Phyllosticta*, especially the homothallic ones. Possible candidates would be species from the *P. musarum* and *P. abieticola* clades, to assess if their mating-type locus is similar to *P. capitalensis*, which would be expected based on our results from the ancestral state reconstruction (Figure 4). Another possibility is to investigate if the maintenance of homothallism provides lifestyle benefits to these species, given the ease of spread and sexual propagation to new locations provided by the unselective mating in homothallic individuals (Nagel et al., 2018; Thynne et al., 2017). Moreover, for the heterothallic species, it would be interesting to assess which advantages arise from the shift to heterothallism, and, especially, how they relate to the evolutionary potential of the pathogenic species, such as *P. citricarpa*. Such possibilities highlight the importance of addressing the *Citrus-Phyllosticta* interactions under an evolutionary and genomic perspective in future, in a whole genome context.

## 4 Conclusions

In summary, our study shows the mating-type locus in *Phyllosticta* as a very dynamic region, with different possible rearrangements that lead to changes in reproduction strategies. Our results demonstrate that although mating-type genes show high sequence variability, the mating-type locus organization is well conserved along *Citrus*-associated *Phyllosticta* species, contrasting evolutionary processes at gene and genomic level. Reconstruction of the ancestral thallism in *Phyllosticta* suggests that homothallism is the ancestral state, with a transition to heterothallism in some of the *Citrus*-associated *Phyllosticta* species mediated by gene losses. Moreover, homothallism in *P. capitalensis* and *P. citrichinaensis* results from different evolutionary events, as the remodeling of the mating-type locus in *P. citrichinaensis* could have happened through a TE-mediated translocation. Idiomorphic-specific versions of the *PIM* gene provide evidence of an expansion of the idiomorphic region in the heterothallic species. As a whole, knowledge on mating-type evolution in *Citrus-*associated *Phyllosticta* species offers new perspectives in the study of *Citrus*-*Phyllosticta* interactions, especially under evolutionary and genomic approaches, and may provide valuable information to guide disease management practices.

## Acknowledgements

Authors are grateful to Doreen Landermann and Mieke Starink-Willemse for technical assistance, to Paulo J. C. dos Santos for the support in generating *P. citribraziliensis* read data, and Primrose Boynton for her feedback on the manuscript.

## Funding

This study was financed in part by the Coordenação de Aperfeiçoamento de Pessoal de Nível Superior - Brasil (CAPES) - Finance Code 001, which provided a PhD scholarship to Desirrê A. L. Petters-Vandresen; it was also supported by the Conselho Nacional de Desenvolvimento Científico e Tecnológico (CNPq) with the INCT-Citros grant 465440/2014-2 and the research grant 424738/2016-3 to Chirlei Glienke, and by FAPESP with the grant 2014/50880-0, allowing the generation of some of the *Phyllosticta* genomes used in this study.

## Supplementary Figures Captions

**Figure S1 – Pairwise comparison of mating-type and flaking genes between *Phyllosticta* species, including all strains evaluated in this study**. The black horizontal lines represent nucleotide genomic sequences, and colour-coded arrows represent coding sequences in forward (right-oriented arrow) or reverse (left-oriented arrow) strand. Boxes between the genomic sequences represent the pairwise similarity based on BLASTn analyses, coded from red (most similar) to blue (least similar). Gene abbreviations: *SAICARsyn*: Phosphoribosylaminoimidazole-succinocarboxamide Synthetase; *MCP*: Mitochondrial carrier protein; *CIA30*: Complex I intermediate-associated protein 30; *APC5*: Anaphase-promoting complex subunit 5; *CoxVIa*: Cytochrome c oxidase subunit Via, *APN2*: DNA lyase; *PIM*: putative integral membrane protein; *40S S9*: 40S ribosomal protein S9. Phylogenetical relationships between *Phyllosticta* species, based in a multilocus dataset of the internal transcribed spacers and intervening 5.8S nrDNA (*ITS*), translation elongation factor 1-alpha (*TEF1*), actin (*ACT*), and glyceraldehyde-3-phosphate dehydrogenase (*GAPDH*) sequences are indicated by the cladogram on the left. ML bootstrap support values above 95% and Bayesian posterior probability values (PP) above 0.9 indicated with an asterisk (*) to the left of the nodes. Scale bar indicates the number of substitutions per nucleotide.

**Figure S2 – Pairwise comparison between mating-type genes and flanking intergenic regions showing homology of *MAT1-1-8* gene fragments in *MAT1-2* strains of *Phyllosticta citriasiana, P. citricarpa* and *P. paracitricarpa***. The black horizontal lines represent nucleotide genomic sequences, and colour-coded arrows represent coding sequences in forward (right-oriented arrow) or reverse (left-oriented arrow) strand. Boxes between the genomic sequences represent the pairwise similarity based on BLASTn analyses, coded from red (most similar) to blue (least similar).

**Figure S3 - Bayesian Inference tree of *SAICARsyn* (a) nucleotide (b) and amino acid sequences of *Phyllosticta* species**. ML bootstrap support values above 95% and Bayesian posterior probability values (PP) above 0.9 indicated with an asterisk (*) to the left of the nodes. Both trees are rooted to *Diplodia seriata*. Scale bar indicates the number of substitutions per nucleotide.

**Figure S4 - Bayesian Inference tree of *MCP* (a) nucleotide (b) and amino acid sequences of *Phyllosticta* species**. ML bootstrap support values above 95% and Bayesian posterior probability values (PP) above 0.9 indicated with an asterisk (*) to the left of the nodes. Both trees are rooted to *Diplodia seriata*. Scale bar indicates the number of substitutions per nucleotide.

**Figure S5 - Bayesian Inference tree of *CIA30* (a) nucleotide (b) and amino acid sequences of *Phyllosticta* species**. ML bootstrap support values above 95% and Bayesian posterior probability values (PP) above 0.9 indicated with an asterisk (*) to the left of the nodes. Both trees are rooted to *Diplodia seriata*. Scale bar indicates the number of substitutions per nucleotide.

**Figure S6 - Bayesian Inference tree of *APC5* (a) nucleotide (b) and amino acid sequences of *Phyllosticta* species**. ML bootstrap support values above 95% and Bayesian posterior probability values (PP) above 0.9 indicated with an asterisk (*) to the left of the nodes. Both trees are rooted to *Diplodia seriata*. Scale bar indicates the number of substitutions per nucleotide.

**Figure S7 - Bayesian Inference tree of *CoxVIa* (a) nucleotide (b) and amino acid sequences of *Phyllosticta* species**. ML bootstrap support values above 95% and Bayesian posterior probability values (PP) above 0.9 indicated with an asterisk (*) to the left of the nodes. Both trees are rooted to *Diplodia seriata*. Scale bar indicates the number of substitutions per nucleotide.

**Figure S8 - Bayesian Inference tree of *APN2* (a) nucleotide (b) and amino acid sequences of *Phyllosticta* species**. ML bootstrap support values above 95% and Bayesian posterior probability values (PP) above 0.9 indicated with an asterisk (*) to the left of the nodes. Both trees are rooted to *Diplodia seriata*. Scale bar indicates the number of substitutions per nucleotide.

**Figure S9 - Bayesian Inference tree of *MAT1-1-1* (a) nucleotide (b) and amino acid sequences of**

***Phyllosticta* species**. ML bootstrap support values above 95% and Bayesian posterior probability values (PP) above 0.9 indicated with an asterisk (*) to the left of the nodes. Both trees are rooted to *Diplodia seriata*. Scale bar indicates the number of substitutions per nucleotide.

**Figure S10 - Bayesian Inference tree of *MAT1-1-8* (a) nucleotide (b) and amino acid sequences of *Phyllosticta* species**. ML bootstrap support values above 95% and Bayesian posterior probability values (PP) above 0.9 indicated with an asterisk (*) to the left of the nodes. Both trees are rooted to *Diplodia seriata*. Scale bar indicates the number of substitutions per nucleotide.

**Figure S11 - Bayesian Inference tree of *MAT1-2-1* (a) nucleotide (b) and amino acid sequences of *Phyllosticta* species**. ML bootstrap support values above 95% and Bayesian posterior probability values (PP) above 0.9 indicated with an asterisk (*) to the left of the nodes. Both trees are rooted to *Diplodia seriata*. Scale bar indicates the number of substitutions per nucleotide.

**Figure S12 - Bayesian Inference tree of *MAT1-2-5* (a) nucleotide (b) and amino acid sequences of *Phyllosticta* species**. ML bootstrap support values above 95% and Bayesian posterior probability values (PP) above 0.9 indicated with an asterisk (*) to the left of the nodes. Both trees are rooted to *Diplodia seriata*. Scale bar indicates the number of substitutions per nucleotide.

**Figure S13 - Bayesian Inference tree of *40S S9* (a) nucleotide (b) and amino acid sequences of *Phyllosticta* species**. ML bootstrap support values above 95% and Bayesian posterior probability values (PP) above 0.9 indicated with an asterisk (*) to the left of the nodes. Both trees are rooted to *Diplodia seriata*. Scale bar indicates the number of substitutions per nucleotide.

**Figure S14 - Bayesian Inference tree of PIM (a) nucleotide (b) and amino acid sequences of *Phyllosticta* species**. ML bootstrap support values above 95% and Bayesian posterior probability values (PP) above 0.9 indicated with an asterisk (*) to the left of the nodes. Both trees are rooted to *Diplodia seriata*. Scale bar indicates the number of substitutions per nucleotide.

## Notes

### Competing Interest Statement

The authors have declared no competing interest.

