## Supplemental Table 1 for "Mating-type locus rearrangement leads to shift from homothallism to heterothallism in *Citrus*-associated *Phyllosticta* species"

Table S1 – Assembly statistics

|  | ***P. capitalensis* LGMF01** | ***P. citricarpa* LGMF06** | ***P. citribraziliensis* LGMF08** |
| --- | --- | --- | --- |
| Reads size (Raw reads) | 101 bp | 101 bp | 250bp |
| Reads size (Trimmed reads) | 50 – 101 bp | 50 – 101 bp | 230 – 250 bp |
| Reads count (Raw reads) | 148,831,020 | 179,880,616 | 248,840,034 |
| Reads count (Trimmed reads) | 66,551,346 | 51,843,994 | 97,631,706 |
| Coverage* (Raw reads) | 461× | 567× | 2006× |
| Coverage (Trimmed reads) | 206× | 163× | 787× |
| Contigs > 0 bp | 2322 | 33409 | 563 |
| Contigs > 1000 bp | 104 | 245 | 422 |
| Contigs > 5000 bp | 44 | 165 | 249 |
| Contigs > 10 kb | 40 | 153 | 200 |
| Total lenght | 32,606,250 | 32,021,677 | 31,002,620 |
| N50 | 1,366,738 | 421,474 | 263,424 |
| L50 | 9 | 28 | 40 |
| GC (%) (ignoring Ns) | 54.55% | 52.83% | 54.34% |

* Coverage estimated by: size of the reads multiplied by the number of reads, divided by the genome size
