## Supplemental Table 2 for "Mating-type locus rearrangement leads to shift from homothallism to heterothallism in *Citrus*-associated *Phyllosticta* species"

Table S2 – Primer names, sequences, annealing temperature and amplicon sizes for the primers designed to confirm the presence of the transposable element in the mating-type locus of *Phyllosticta citrichinaensis*

| Primer name | Primer Sequence (5’-3’) | Primer annealing positions in  *P. citrichinaensis*^a^ | Amplicon size | Description / Target | Source |
| --- | --- | --- | --- | --- | --- |
| MAT111degF2 | GCTCTCAACTCTTTCATGGC | 14973 | 5,021 bp | *Phyllosticta* spp. *MAT1-1-1* specific | (Guarnaccia et al., 2019) |
| TE-Citrichina-R1 | CCTCTAAGGTAGACCC | 19994 |  | LTR-*Copia* element in *P. citrichinaensis* *MAT* locus | This study |
| TE-Citrichina-F1 | GGGTCTACCTTAGAGG | 19979 | 6,474 bp | LTR-*Copia* element in *P. citrichinaensis* *MAT* locus | This study |
| PIM-Citrichina-R1 | GTTCCACGAGACTCTCC | 26453 |  | LTR-*Copia* element in *P. citrichinaensis* *MAT* locus | This study |

^a^Position in relation to the first base of the *MAT* locus sequence of *P. citrichinaensis* provided in Supplementary Material
