## Supplemental Table 3 for "Mating-type locus rearrangement leads to shift from homothallism to heterothallism in *Citrus*-associated *Phyllosticta* species"

Table S3 – Host/substrate, geographical origin, collection codes and GenBank accession numbers of *Phyllosticta* spp. and *Botryosphaeriales* reference strains used for phylogenetic analyses

| **Species** | **Host/Substrate** | **Origin** | **Collection^a^** | **GenBank accession^b^** | | | | **References** |
| --- | --- | --- | --- | --- | --- | --- | --- | --- |
|  |  |  |  | **ITS** | ***ACT*** | ***GAPDH*** | ***TEF1*** |  |
| *P. abieticola* | *Abies concolor* | Canada | **CBS 112067** | KF170306 | KF289238 | - | - | (Wikee et al., 2013) |
| *P. ampelicida* | *Vitis riparia* | USA | **ATCC 200578** | KC193586 | KC193581 | KC193584 | - | (Zhang et al., 2013) |
| *P. capitalensis* | *Stanhopea graveolens* | Brazil | **CBS 128856** | JF261465 | JF343647 | JF343776 | JF261507 | (Glienke et al., 2011) |
| *P. cavendishii* | *Musa* sp. cv. Formosana (AAA) | Taiwan | **BRIP 55420** | JQ743562 | - | - | - | (Wong et al., 2012) |
| *P. citriasiana* | *Citrus maxima* | Thailand | **CBS 120486** | FJ538360 | FJ538476 | JF343686 | FJ538418 | (Glienke et al., 2011; Wulandari et al., 2009) |
| *P. citribraziliensis* | *Citrus* sp. | Brazil | **CBS 100098** | FJ538352 | FJ538468 | JF343691 | FJ538410 | (Glienke et al., 2011; Wulandari et al., 2009) |
|  | *Citrus* sp. | Brazil | LGMF08 | JF261435 | JF343617 | JF343617 | JF261477 | (Glienke et al., 2011) |
| *P. citricarpa* | *Citrus limon* | Australia | **CBS 127454** | JF343583 | JF343667 | JF343771 | JF343604 | (Glienke et al., 2011) |
|  | *Citrus sinensis* | Brazil | LGMF06 | JF261431 | JF343613 | JF343676 | JF261473 | (Glienke et al., 2011) |
|  | *Citrus sinensis* | Malta | CPC 27913 | KY855614 | KY855669 | KY855724 | KY855943 | (Guarnaccia et al., 2017) |
| *P. citrichinaensis* | *Citrus maxima* | China | **CBS 130529** | JN791597 | JN791526 | KY855734 | JN791452 | (Guarnaccia et al., 2017; Wang et al., 2012) |
| *P. citrimaxima* | *Citrus maxima* | Thailand | **CBS 136059** | KF170304 | KF289300 | KF289157 | KF289222 | (Wikee et al., 2013) |
| *P. maculata* | *Musa* sp. cv. Goly-goly pot-pot (ABB) | Australia | **CBS 132581** | JQ743570 | - | - | - | (Wong et al., 2012) |
| *P. musarum* | Hill banana (AAB) | India | **BRIP 55434** | JQ743584 | - | - | - | (Wong et al., 2012) |
| *P. paracapitalensis* | *Citrus × floridana* | Italy | **CBS 141353** | KY855622 | KY855677 | KY855735 | KY855951 | (Guarnaccia et al., 2017) |
| *P. paracitricarpa* | *Citrus limon* | Greece | **CBS 141357** | KY855635 | KY855690 | KY855748 | KY855964 | (Guarnaccia et al., 2017) |
| *P. rubella* | *Acer rubrum* | USA | **CBS 111635** | KF206171 | KF289233 | KF289129 | KF289198 | (Wikee et al., 2013) |
| *Diplodia corticola* | *Quercus suber* | Portugal | **CBS 112549** | AY259100 | MG015715 | * | AY573227 | (Alves et al., 2004; Lopes et al., 2018; Phillips et al., 2005) |
| *Neofusicoccum parvum* | *Syzygium cordatum* | South Africa | **CBS 123649** | KX464213 | * | * | KX464743 | (Yang et al., 2017) |

^a^Type strains included in analyses are indicated in **bold.** Culture collection abbreviations: ATCC = American Type Culture Collection, Virginia, USA; BRIP = Queensland Plant Pathology Herbarium, Brisbane, Australia; CBS = Westerdijk Fungal Biodiversity Institute, Utrecht, The Netherlands; CGMCC = Chinese General Microbiological Culture Collection Center, Beijing, China; CPC = Culture collection of Pedro W. Crous, held at Westerdijk Fungal Biodiversity Institute; LGMF = Laboratório de Bioprospecção e Genética Molecular de Microrganismos, Universidade Federal do Paraná, Paraná, Brazil;

^b^GenBank - ITS*:* internal transcribed spacers and intervening 5.8S nrDNA*; ACT:* partial actin gene*; GAPDH:* partial glyceraldehyde-3-phosphate dehydrogenase gene*; TEF1:* partial translation elongation factor 1-alpha gene*.* -: Information or sequence not available; *: Sequence retrieved from the whole genome assembly of the strain through BLAST.
