## Supplemental Table 4 for "Mating-type locus rearrangement leads to shift from homothallism to heterothallism in *Citrus*-associated *Phyllosticta* species"

Table S4 – Characteristics of the gene partitions used in phylogenetic analyses

| **Dataset** | **Locus^a^** | **Number of sites** | | | | **Evolutionary model^b^** |
| --- | --- | --- | --- | --- | --- | --- |
|  |  | **Total** | **Constant** | **Variable** | **Parsimony informative** |  |
| *Citrus*-associated *Phyllosticta* species for synteny figure of type-strains (nucleotides) | ITS | 600 | 512 | 80 | 45 | SYM+G |
|  | *TEF1* | 301 | 241 | 50 | 15 | GTR |
|  | *ACT* | 274 | 236 | 31 | 12 | HKY+G |
|  | *GAPDH* | 651 | 597 | 53 | 25 | GTR+I |
| *Citrus*-associated *Phyllosticta* species for synteny figure of all strains (nucleotides) | ITS | 596 | 509 | 87 | 82 | SYM+G |
|  | *TEF1* | 261 | 214 | 47 | 36 | GTR |
|  | *ACT* | 269 | 226 | 39 | 36 | HKY |
|  | *GAPDH* | 651 | 598 | 53 | 49 | GTR+G |
| *Phyllosticta* genus for ancestral reconstruction (nucleotides) | ITS | 636 | 442 | 183 | 125 | GTR+G |
|  | *TEF1* | 326 | 154 | 144 | 92 | GTR+I |
|  | *ACT* | 301 | 196 | 97 | 59 | HKY+G |
|  | *GAPDH* | 655 | 514 | 138 | 76 | GTR+I |
| *Citrus*-associated *Phyllosticta* species and Botryosphaeriaceae members (nucleotides) | *SAICARsyn* | 1080 | 650 | 320 | 190 | HKY+G |
|  | *MCP* | 1167 | 797 | 360 | 250 | HKY+I |
|  | *CIA30* | 783 | 514 | 246 | 139 | HKY+G |
|  | *APC5* | 2838 | 1554 | 1112 | 721 | HKY+I+G |
|  | *CoxVIa* | 407 | 301 | 97 | 36 | HKY+G |
|  | *APN2* | 2220 | 1258 | 817 | 513 | GTR+I |
|  | *MAT1-1-8* | 1441 | 664 | 555 | 220 | HKY+I |
|  | *MAT1-1-1* | 2087 | 1121 | 831 | 313 | HKY+I |
|  | *MAT1-2-1* | 1524 | 757 | 639 | 156 | GTR+I |
|  | *MAT1-2-5* | 1204 | 594 | 442 | 72 | HKY+G |
|  | *PIM* | 4333 | 1217 | 1490 | 1142 | HKY+I+G |
|  | *40S S9* | 582 | 503 | 76 | 31 | GTR+I |
| *Phyllosticta* + Botryosphaeriaceae members (amino acids) | *SAICARsyn* | 350 | 206 | 110 | 68 | WAG+G |
|  | *MCP* | 386 | 314 | 71 | 37 | JTT+I |
|  | *CIA30* | 256 | 158 | 93 | 41 | JTT+G |
|  | *APC5* | 931 | 456 | 430 | 244 | JTT+G |
|  | *CoxVIa* | 136 | 93 | 38 | 19 | JTT+I |
|  | *APN2* | 727 | 411 | 289 | 157 | JTT+G |
|  | *MAT1-1-8* | 474 | 146 | 266 | 70 | LG+G |
|  | *MAT1-1-1* | 688 | 249 | 389 | 106 | JTT+I+G |
|  | *MAT1-2-1* | 493 | 160 | 296 | 33 | JTT+I |
|  | *MAT1-2-5* | 390 | 93 | 254 | 30 | JTT+G |
|  | *PIM* | 1418 | 324 | 568 | 443 | JTT+G |
|  | *40S S9* | 193 | 188 | 4 | 2 | LG |

^a^Locus abbreviations: ITS: internal transcribed spacers and intervening 5.8S nrDNA; *tef1*: Translation elongation factor 1-alpha; *act*: actin; *gapdh*: glyceraldehyde-3-phosphate dehydrogenase; *SAICARsyn*: Phosphoribosylaminoimidazolesuccinocarboxamide Synthetase; *MCP*: Mitochondrial carrier protein; *CIA30*: Complex I intermediate-associated protein 30; *APC5*: Anaphase-promoting complex subunit 5; *CoxVIa*: Cytochrome c oxidase subunit VIa, *APN2*: DNA lyase; *PIM*: putative integral membrane protein; *40S S9*: 40S ribosomal protein S9.

^b^Evolutionary model: G: Gamma distributed rate variation among sites; GTR: Generalised time-reversible model; HKY: Hasegawa, Kishino and Yano model; I: Proportion of invariable sites; JTT: Jones-Taylor-Thornton model; LG: Le and Gascuel model; SYM: Symmetrical model; WAG: Whelan And Goldman model.
