## Supplemental Table 5 for "Mating-type locus rearrangement leads to shift from homothallism to heterothallism in *Citrus*-associated *Phyllosticta* species"

Table S5a – Coding domain sequence sizes, intron sizes and start positions of mating type genes of *Citrus*-associated *Phyllosticta* species

|  | *MAT 1-1-1* | | | *MAT 1-1-8* | | | *MAT 1-2-1* | | | *MAT 1-2-5* | | |
| --- | --- | --- | --- | --- | --- | --- | --- | --- | --- | --- | --- | --- |
| Species | CDS size | Intron size | Intron start position^a^ | CDS size | Intron sizes | Intron start positions^a^ | CDS size | Intron sizes | Intron start positions^a^ | CDS size | Intron sizes | Intron start positions^a^ |
| *P. capitalensis* | 1863 | 51 | 440 | 1089 | 48, 57, 149, 138 | 155, 715, 805, 1047 | 1389 | 49, 51 | 333, 717 | 1041 | 49, 66 | 609, 936 |
| *P. citriasiana* | 1881 | 52 | 437 | 1098 | 268, 52, 133, 99 | 146, 926, 1011, 1237 | 1359 | 53, 51 | 324, 691 | 996 | 48, 66 | 579, 905 |
| *P. citribraziliensis* | 1869 | 51 | 440 | 1077 | 47, 48, 111, 139 | 117, 676, 757, 961 | - | - | - | - | - | - |
| *P. citricarpa* | 1890 | 51 | 437 | 1098 | 267, 49, 131, 96 | 146, 925, 1007, 1231 | 1359 | 53, 51 | 324, 691 | 996 | 48,66 | 579, 905 |
| *P. citrichinaensis* | 1848 | 51 | 428 | 1059 | 51, 48, 114, 113 | 116, 679, 760, 967 | 1356 | 51, 50 | 324, 686 | 1008 | 48, 65 | 579, 905 |
| *P. paracitricarpa* | - | - | - | - | - | - | 1359 | 53, 51 | 324, 691 | 996 | 48, 66 | 579, 905 |

^a^Intron start position measured in number of base pairs from the first base of the gene.

Table S5b – Coding domain sequence sizes, intron sizes and start positions of mating type adjacent genes of *Citrus*-associated *Phyllosticta* species

|  | *SAICARsyn*^a^ | | | *MCP*^a^ | | | *CIA30*^a^ | | | *APC5*^a^ | | |
| --- | --- | --- | --- | --- | --- | --- | --- | --- | --- | --- | --- | --- |
| Species | CDS size | Intron sizes | Intron start positions^b^ | CDS size | Intron sizes | Intron start positions^b^ | CDS size | Intron size | Intron start position^b^ | CDS size | Intron sizes | Intron start positions^b^ |
| *P. capitalensis* | 927 | 276, 114, 78, 97 | 160, 806, 987 | 1152 | 52, 47 | 186, 986 | 708 | 55 | 83 | 2556 | 49, 45, 46, 49, 47, 70, 134 | 686, 829, 1286, 1456, 1612, 1796, 2546 |
| *P. citriasiana* | 927 | 160, 85, 84, 77 | 157, 696, 848 | 1152 | 56, 49 | 186, 990 | 750 | 56 | 83 | 2430 | 52, 46, 46, 48, 52, 87, 82, 138 | 683, 829, 1287, 1457, 1612, 1801, 2315, 2647 |
| *P. citribraziliensis* | 924 | 140, 82, 62, 89 | 154, 673, 822 | 1152 | 46, 44 | 186, 980 | 750 | 50 | 83 | 2481 | 45, 46, 45, 46, 49, 97, 66, 121 | 680, 819, 1277, 1446, 1599, 1785, 2309, 2607 |
| *P. citricarpa* | 927 | 160, 84, 84, 77 | 157, 696, 847 | 1152 | 56, 49 | 186, 990 | 750 | 56 | 83 | 2430 | 52, 45, 46, 48, 52, 87, 82, 132 | 683, 829, 1286, 1456, 1611, 1800, 2314, 2646 |
| *P. citrichinaensis* | 924 | 136, 97, 102, 82 | 154, 669, 833 | 1152 | 46, 44 | 186, 980 | 750 | 50 | 83 | 2478 | 46, 46, 44, 46, 50, 104, 68, 91 | 680, 820, 1278, 1446, 1599, 1786, 2320, 2614 |
| *P. paracitricarpa* | 927 | 160, 84, 94, 77 | 157, 696, 847 | 1152 | 56, 48 | 186, 990 | 750 | 56 | 83 | 2439 | 52, 45, 46, 48, 52, 87, 82, 132 | 683, 829, 1286, 1456, 1611, 1800, 2314, 2655 |

^a^Gene abbreviations: *SAICARsyn*: Phosphoribosylaminoimidazolesuccinocarboxamide Synthetase; *MCP*: Mitochondrial carrier protein; *CIA30*: Complex I intermediate-associated protein 30; *APC5*: Anaphase-promoting complex subunit 5.

^b^Intron start position measured in number of base pairs from the first base of the gene.

Table S5c: Coding domain sequence size, intron size and start position of mating type adjacent genes of *Citrus*-associated *Phyllosticta* species

|  | *COX VIa*^a^ | | | *DNA lyase*^a^ | | | *40S S9*^a^ | | |
| --- | --- | --- | --- | --- | --- | --- | --- | --- | --- |
| Species | CDS size | Intron sizes | Intron start positions^b^ | CDS size | Intron sizes | Intron start positions^b^ | CDS size | Intron sizes | Intron start positions^b^ |
| *P. capitalensis* | 393 | 62, 185, 210 | 164, 251, 596 | 1968 | 46, 45, 180 | 224, 980, 1959 | 576 | 94, 54, 52 | 66, 259, 547 |
| *P. citriasiana* | 393 | 74, 179, 189 | 164, 263, 602 | 1998 | 45, 49, 221 | 221, 976, 1992 | 579 | 94, 86, 88 | 66, 259, 579 |
| *P. citribraziliensis* | 393 | 69, 164, 158 | 164, 258, 582 | 1995 | 49, 46, 201 | 221, 980, 1990 | 579 | 91, 55, 64 | 66, 256, 545 |
| *P. citricarpa* | 393 | 74, 178, 182 | 164, 263, 601 | 1998 | 45, 49, 221 | 221, 976, 1992 | 579 | 94, 87, 88 | 66, 259, 580 |
| *P. citrichinaensis* | 393 | 71, 169, 145 | 164, 260, 589 | 2004 | 45, 47, 193 | 221, 976, 1996 | 579 | 91, 55, 65 | 66, 256, 545 |
| *P. paracitricarpa* | 393 | 74, 178, 182 | 164, 263, 601 | 1998 | 45, 49, 221 | 221, 976, 1992 | 579 | 94, 87, 88 | 66, 259, 580 |

^a^Gene abbreviations: *CoxVIa*: Cytochrome c oxidase subunit VIa, *APN2*: DNA lyase; *40S S9:* 40S ribosomal protein S9.

^b^Intron start position measured in number of base pairs from the first base of the gene

Table S5d: Coding domain sequence size, intron size and start position of mating type adjacent genes of *Citrus*-associated *Phyllosticta* species

|  | *PIM*^a^ | | |
| --- | --- | --- | --- |
| Species | CDS size | Intron size | Intron start position^b^ |
| *P. capitalensis* | 2100 | 51 | 465 |
| *P. citriasiana* (*MAT1-1* idiomorph) | 2529 | 49 | 459 |
| *P. citriasiana* (*MAT1-2* idiomorph) | 2157 | 51 | 480 |
| *P. citribraziliensis* | 2544 | 52 | 459 |
| *P. citricarpa* (*MAT1-1* idiomorph) | 2529 | 49 | 459 |
| *P. citricarpa* (*MAT1-2* idiomorph) | 2157 | 51 | 480 |
| *P. citrichinaensis* | 2163 | 51 | 483 |
| *P. paracitricarpa* | 2172 | 51 | 480 |

^a^Gene abbreviations: *PIM*: putative integral membrane protein.

^b^Intron start position measured in number of base pairs from the first base of the gene
